# Reference-free single-vesicle profiling of small extracellular vesicles from liquid biopsies with the PICO assay

**DOI:** 10.64898/2026.02.27.707718

**Authors:** John Atanga, Pablo Sánchez-Martín, Siobhan King, Tobias Gross, Irina Nazarenko

## Abstract

Small extracellular vesicles (sEV) are membrane-enclosed nanoparticles found in body fluids that carry molecular cargo from their cells of origin. Their stability and disease-associated molecular signatures make them promising targets for the development of non-, or minimally invasive liquid biopsies, yet scalable approaches enabling single-vesicle quantification of sEV while resolving their heterogeneity remain limited. Here, we present PICO (Protein Interaction Coupling), a reference-free quantitative assay adapted for sensitive multiplex profiling of individual intact vesicles. PICO detects vesicle markers by requiring colocalization of two or more copies of the same protein or of distinct proteins (e.g., CD9 or CD9/CD63) on individual vesicles, using DNA-barcoded antibodies and digital PCR (dPCR) for quantitative readout. We demonstrate that this unique architecture of the assay provides high specificity by distinguishing EV-bound proteins from soluble counterparts, and can be adapted to target either surface-exposed or intravesicular biomarkers. PICO requires minimal sample input (1 µl) and no specialized instrumentation beyond standard digital PCR. In a head-to-head comparison with nano-flow cytometry, PICO achieved a comparable limit of detection for sEV subpopulations. Profiling sEV isolates from blood for canonical markers (CD9, CD63 and CD81) and HER2 demonstrates precise, high-resolution quantification of sEV subpopulations in complex clinical samples and supports integration of scalable single-EV analysis into research and diagnostic workflows.

## Main

Liquid biopsies offer non-, and minimally invasive access to molecular information from tumors and host tissues. It is complementary to tissue biopsy as they can support early cancer detection, longitudinal monitoring of therapy response, identification of minimal residual disease and support the implementation of companion diagnostics, thereby enabling more adaptive and personalized therapeutic strategies^1,2^. In clinical oncology, liquid biopsies are particularly valuable when tumor tissue is inaccessible, insufficient or cannot be repeatedly sampled, and they allow real-time assessment of disease dynamics during treatment. Currently, analysis of circulating cell-free DNA (cfDNA) represents the most established liquid biopsy modality and is integrated into clinical guidelines across multiple cancer entities^3–5^. However, cfDNA provides an intrinsically tumor-centric and incomplete view of disease biology. cfDNA is predominantly released through cell-death processes, particularly apoptosis and necrosis, and is therefore enriched for DNA fragments derived from dying or highly proliferative tumor cells^6–9^. As a consequence, cfDNA may underrepresent dormant tumor cell populations and provides limited insight into systemic responses to therapy, including immune activation, stromal remodeling and organ-wide stress responses. Thus, while cfDNA is highly informative for genetic alterations within tumors, it does not capture the full biological complexity of cancer as a systemic disease. Extracellular vesicles (EV) represent a complementary and potentially synergistic class of liquid biopsy biomarkers. EV are lipid bilayer–enclosed particles actively secreted by virtually all cell types and are present in bodily fluids such as blood, urine and cerebrospinal fluid^10–12^. Both on their surface and within their lumen, EV carry proteins, lipids and nucleic acids that reflect the molecular state of their cells of origin, enabling them to report on dynamic cellular processes in living tissue^13^. Unlike cfDNA, EV are predominantly released by viable cells and therefore circulate as heterogeneous populations of vesicles from diverse cellular sources, including tumor, stroma and immune cells^14,15^. Because EVs derive from both tumor and non-malignant cells, EV-based liquid biopsies can report on tumor-associated signals alongside therapy-related systemic processes such as immune response, providing a more comprehensive view of the patient’s physiological state during treatment. Despite their promise, translation of EV-based research findings into clinical practice is facing significant technical and methodological hurdles^16–18^. EV populations in biofluids are highly heterogeneous and comprise vesicles originating from diverse cell types, making population-level discrimination by cellular origin and functional state essential. Currently, most scalable EV analysis approaches rely on bulk measurements that average signals across vesicle populations, thereby obscuring biologically and clinically relevant EV subpopulations^19^. Single-vesicle analysis techniques, including nano-flow cytometry and interferometry-based platforms, have substantially advanced the field and enable multiparametric phenotyping of individual EVs. However, these approaches typically require specialized and costly instrumentation, complex calibration procedures and expert operation, which limit standardization, scalability and routine implementation in clinical and translational settings. As EV populations in biofluids are highly heterogeneous and derive from multiple cellular sources, analysis of complex EV mixtures requires quantitative, single-vesicle methods that can reliably assess co-localized biomarkers on individual vesicles. While co-detection of multiple markers is technically feasible with existing platforms^20–22^, there remains a need for approaches that enable such analyses in a broadly accessible, reference-free and easily scalable format. Consequently, methods that combine quantitative single-vesicle resolution with multiplexed biomarker co-localization, without reliance on specialized infrastructure, are essential for systematic dissection of EV subpopulations in clinical samples^23,24^. Here, we extend the application of PICO, a digital PCR immunoassay enabling absolute protein quantification^25^ to the detection and quantification of sEV. This approach enables sensitive, quantitative single-vesicle analysis with performance comparable to state-of-the-art technologies, while requiring no specialized instrumentation or reference calibration materials, incorporating an explicit quality-control step to preserve and verify EV integrity, and providing a practical route toward scalable implementation in translational and clinical workflows.

### The fundamental principle of PICO assay for sEV detection

In the previous work^25^, we established the conceptual and statistical framework of the PICO assay for reference-free absolute protein quantification. In brief, pairs of DNA-labeled antibodies bind simultaneously to a target molecule, forming a ternary complex termed a *couplex* (Fig. 1a). After binding equilibrium is reached, the reaction mixture is diluted and compartmentalized in a digital PCR (dPCR) device, where the antibody-linked DNA labels are amplified and detected using fluorescent probes. This enables classification of partitions as negative, single-positive or double-positive for the respective antibody labels^25^. For soluble targets, four binding states are possible: no antibody binding, binding of either antibody alone, or binding of both antibodies (Fig. 1b). Only the latter results in formation of a true *couplex*. By analyzing the distribution of empty, single-positive and double-positive partitions and applying a compartmental *de-couplexing* statistical model, the number of true *couplexes* can be inferred while accounting for random co-partitioning of antibodies. Under saturating antibody conditions, the equilibrium of these states shifts toward *couplex* formation, enabling absolute target quantification without the need for reference standards^25^. When applied to sEV, the binding landscape differs fundamentally (Fig. 1c,d). Individual vesicles typically present multiple copies of a given surface marker, giving rise to numerous possible antibody–EV binding configurations. Because detection of an EV requires binding of at least one of each antibody in the DNA-labeled pair, the majority of productive binding states result in detectable *couplexes.* Moreover, the presence of multiple marker copies per vesicle increases the probability of antibody binding. Consequently, under saturating antibody conditions, the reaction equilibrium strongly favors *couplex* formation, such that the number of detected *couplexes* directly reflects the concentration of intact sEV in the sample (Fig. 1d).

**Figure 1.**
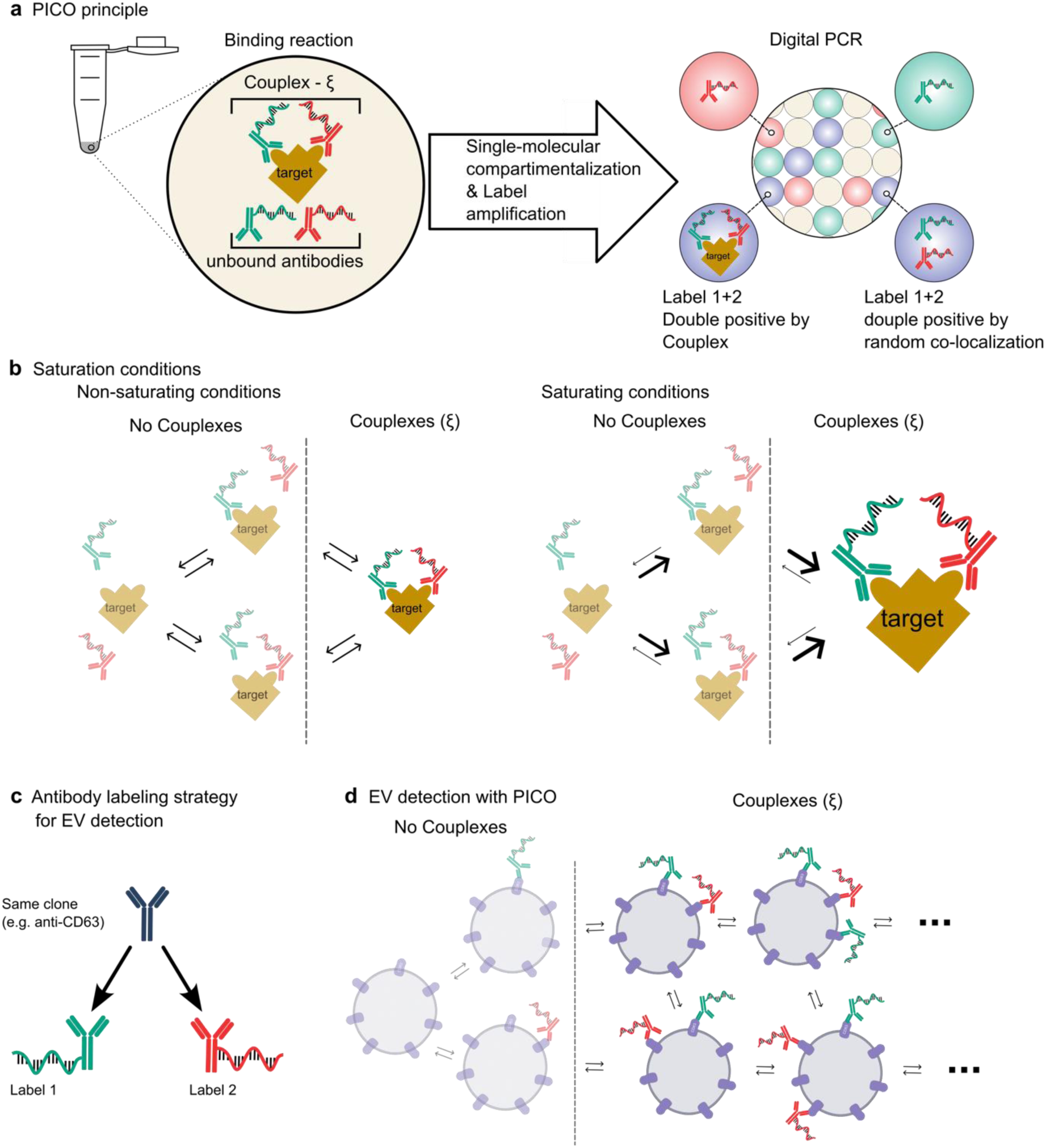
Fundamental principle of the PICO assay for single-step absolute quantification of sEV. **a**, The core PICO assay mechanism. DNA-labeled antibodies bind to target respective epitopes on a target molecule, forming a ternary complex (a “couplex”). The mixture is diluted, compartmentalized via digital PCR (dPCR), and the DNA barcodes in the antibodies are amplified and detected via fluorescence. **b**, Possible binding states for a soluble, monovalent target. Only the state where both antibodies are bound forms a detectable couplex. At saturating antibody concentrations, the equilibrium is shifted towards couplex formation, enabling absolute quantification without a standard curve. **c**, Antibody labeling strategy for the detection of sEV positive for a single marker. The same antibody clone (e.g. anti-CD63) is split and labeled with two distinct oligonucleotide labels. **d**, Simplified PICO assay for EV quantification. Due to the high avidity and multivalency of vesicles, numerous antibody-binding states are possible. Crucially, any EV that has bound at least one of each type of DNA-labeled antibody forms a detectable couplex. Under saturating antibody conditions, the number of detected couplexes directly corresponds to the concentration of EVs

## Results

### Biological reference material for PICO EV detection assay development

Although biological EV reference materials, including commercially available lyophilized preparations, are available, broadly accepted, fit-for-purpose standards for benchmarking and cross-platform comparability remain limited^19,26^. To support PICO assay development and benchmarking, we established and characterized cell-based model that generates sEV with defined, reproducible canonical EV marker profiles approaching physiological levels, for use as biological reference material and assay control. We employed human fibrosarcoma cell line HT1080 to generate sEV with defined tetraspanin profiles. Parental HT1080 cells produce sEV expressing CD63 and CD81 but lacking CD9, serving as a negative control for CD9 detection. In parallel, an HT1080-CD9 knock-in derivative was generated, producing CD9^+^ sEV while retaining expression of CD63 and CD81. Tetraspanin expression was validated at the cellular level by immunoblotting and flow cytometry. CD9 expression was detected exclusively in the HT1080-CD9 knock-in cells and was absent in parental controls, whereas CD63 and CD81 expression levels were comparable between both models, confirming successful CD9 introduction without perturbation of other canonical EV markers (Extended Data Fig. 1a,b). Cell viability exceeded 95% at the time of harvest for both cell lines, indicating that EV release predominantly reflected secretion from viable cells (Extended Data Fig. 1c). sEV were isolated from conditioned media using a combination of tangential flow filtration (TFF) and size-exclusion chromatography (SEC). SEC fractions 1-3 were enriched for sEV, as indicated by high particle concentrations and low protein content, consistent with minimal contamination by soluble proteins (Extended Data Fig. 1d,e). Immunoreactivity for the canonical sEV markers CD9, CD63 and CD81 in these fractions was confirmed by dot blot analysis (Extended Data Fig. 1f,g). Based on this characterization, fractions 1–3 were selected as biological reference sEV, pooled, concentrated and subjected to comprehensive characterization. Particle size and concentration were assessed by nanoparticle tracking analysis (NTA) and nano-flow cytometry (nFCM), and statistical analyses and data visualization were performed using PHoNUPS, an open-source analysis and visualization tool developed by our team^27^. CD9 knock-in did not significantly alter sEV size distributions but resulted in a modest increase in sEV release from HT1080-CD9 cells compared with parental controls (Fig. 2b-g). Immunoblotting of isolated sEV confirmed CD9 expression in HT1080-CD9 - derived vesicles, with comparable levels of CD63 and CD81 between both sEV populations (Fig. 2h). Cryo-electron microscopy revealed that sEV isolated from both HT1080 and HT1080-CD9 conditioned media exhibited intact lipid bilayers and characteristic vesicular morphology (Fig. 2i), confirming vesicle integrity as a prerequisite for their detection and quantification by the PICO assay.

**Figure 2.**
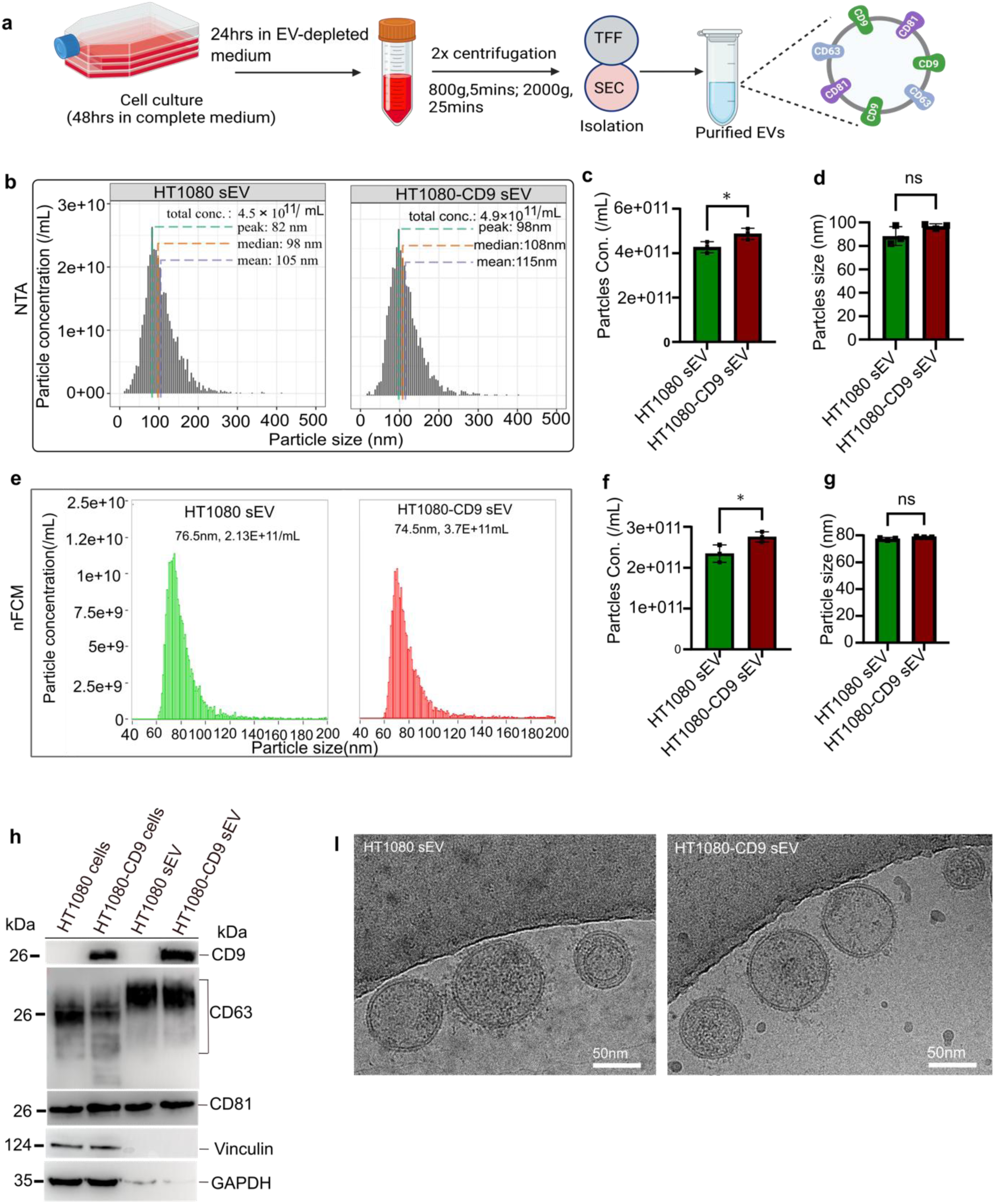
Characterization of biological reference sEV with defined tetraspanin profiles. **a**, Schematic of sEV isolation workflow using tangential flow filtration (TFF) followed by size-exclusion chromatography (SEC). **b, c,** Nanoparticle tracking analysis (NTA) characterization of sEV isolated from parental HT1080 (CD9⁻) and HT1080-CD9 (CD9⁺) cells. Representative particle size distribution profiles **(b)** and quantification of particle concentration **(c)**. Comparison of sEV median size of HT1080 and HT1080-CD9 (**d)** data in **c, d** is presented as mean ± s.d. (n=3 biological replicates); P value was calculated using an unpaired two-tailed t-test. **e-g,** High-resolution single-particle characterization of sEVs by nanoflow cytometry (nFCM). Representative particles size distribution **(e)** and quantification of vesicle concentration **(f)** and size **(g)** for HT1080 and HT1080-CD9 sEV. For the NTA data analysis and visualization of NTA and nano flow cytometry data PHoNUPS open source software was employed^26^. Data in **f** and **g** are presented as mean ± s.d. (n=3 biological replicates); n.s., not significant; P value was calculated using an unpaired two-tailed t-test. **h,** immunoblot analysis of sEVs and cell lysates from HT1080 and HT1080-CD9 cells confirms the presence of CD9 specifically in the knock-in line and comparable expression of CD63 and CD81. Vinculin was used as a negative control for sEV purity, whereas GAPDH served as a loading control. **i,** Representative cryo-electron microscopy (cryo-EM) images of sEV from both cell lines, showing intact vesicles with characteristic morphology and lipid bilayers. Scale bars, 50 nm.

### PICO achieves single-vesicle resolution of canonical tetraspanins

To provide orthogonal validation for PICO-based sEV measurements, canonical EV markers (CD9, CD63 and CD81) were independently assessed on the biological reference sEV by nano-flow cytometry (nFCM) and super-resolution imaging. For nFCM, sEV were stained with monoclonal antibodies against CD9, CD81 and CD63 and analyzed at single-vesicle resolution. Parental HT1080 sEV showed no detectable CD9 signal, whereas subsets were positive for CD81 (49%) and CD63 (18%) (Extended Data Fig. 2a,b). In HT1080-CD9 sEV, the proportions of CD63^+^ and CD81^+^ vesicles were similar to parental controls, while ∼90% of vesicles were CD9^+^ (Fig. 3a-c), corresponding to 1.6 × 10⁷ vesicles/μl (CD9^+^, 90%), 4.9 × 10⁶ vesicles/µl (CD81^+^, 48%) and 2.8 × 10⁶ vesicles/µl (CD63^+^, 17%) (Fig. 3a-c). These markers were next validated by direct stochastic optical reconstruction microscopy (dSTORM). sEV were captured (EV Profiler kit, ONI) and stained with fluorescent antibodies against CD9, CD81 and CD63. Clustering analysis confirmed high CD9 prevalence (90.3%), with subpopulations positive for CD81 (50.6%) and CD63 (20.2%) (Fig. 3d,e), in close agreement with the nFCM-based measurements. To enable PICO-based single-vesicle readout, monoclonal antibodies against CD9, CD81 and CD63 were conjugated to distinct oligonucleotide tags (Fig. 3f). In this configuration, a vesicle is scored as marker-positive when it carries at least one copy of each oligo-tagged antibody of the corresponding pair, resulting in formation of a detectable *couplex* on the vesicle surface. To validate this approach, we incubated serial dilutions of HT1080-CD9 sEV with the corresponding oligo-tagged antibody pairs targeting CD9, CD81 or CD63 (Fig. 3f). An antibody-only control (ABC; no sEV) was included to capture background arising from nonspecific antibody aggregation and amplification artefacts. After normalization and subtraction of ABC values (Methods), we calculated corrected *couplex* counts, representing antibody-pair co-localization on single vesicles, and obtained dilution curves for each marker (Fig. 3g). Inter-assay and biological replicate variability was consistently below 20% (Extended Data Fig. 2c). The limit of detection (LOD) and limit of quantification (LOQ) were 1.15 × 10⁶ sEV/µl and 3.81 × 10⁶ sEV/µl, respectively (Extended Data Fig. 2d). Notably, for the CD9+ population, *couplexes* remained detectable down to 8.26 × 10⁴ sEV/µl, indicating that assay sensitivity depends on the abundance of the corresponding EV subpopulation in the sample. To enable absolute quantification of marker-positive sEV by PICO, it is essential that antibody binding is performed under saturating conditions, such that each vesicle carrying the target marker is bound by the antibody pair and yields a detectable *couplex*. To verify that the antibody concentration used for sEV analysis (5 × 10⁻¹⁰ M) achieves saturation, we incubated a fixed amount of HT1080-CD9 sEV with increasing concentrations of CD9-specific antibodies. As expected, once saturation is reached, further increase in antibody concentration does not result in higher *couplex* counts, indicating that all available vesicles are already bound. Indeed, *couplex* counts reached a plateau at 5 × 10⁻¹⁰ M antibody concentration, confirming saturating binding conditions (Extended Data Fig. 2h), consistent with our previous observations for soluble protein targets^24^. Under these conditions, absolute sEV concentrations were determined from the linear range of the dilution series, in which decreasing sample concentration resulted in proportional reductions in *couplex* counts, as described previously²⁵. Using this approach, PICO quantified 1.68 × 10⁷ sEVs/µl for CD9⁺, 4.4 × 10⁶ sEVs/µl for CD81⁺ and 2.1 × 10⁶ sEVs/µl for CD63⁺ vesicles (Fig. 3h). These values showed strong agreement with independent measurements by nano-flow cytometry (Fig. 3j-l). When PICO-based absolute concentrations were compared across dilution series with nanoparticle tracking analysis (NTA), a linear correlation with a slope of 1.1 was observed, demonstrating robust quantification across a range of sEV concentrations (Extended Data Fig. 2d). HT1080-derived sEV, which lack CD9 expression, showed no significant CD9 *couplex* signal compared to the antibody-only control, confirming assay specificity (Extended Data Fig. 2e). Finally, to assess whether PICO selectively detects intact vesicles, sEV were lysed prior to analysis. Under these conditions, no *couplexes* were detected in the single-marker PICO assay, indicating that PICO specifically quantifies intact sEV and does not detect solubilized surface markers (Extended Data Fig. 2f,g). Together, these results validate PICO as a robust approach for absolute quantification of intact single vesicles based on surface marker expression, implemented using digital PCR.

**Figure 3.**
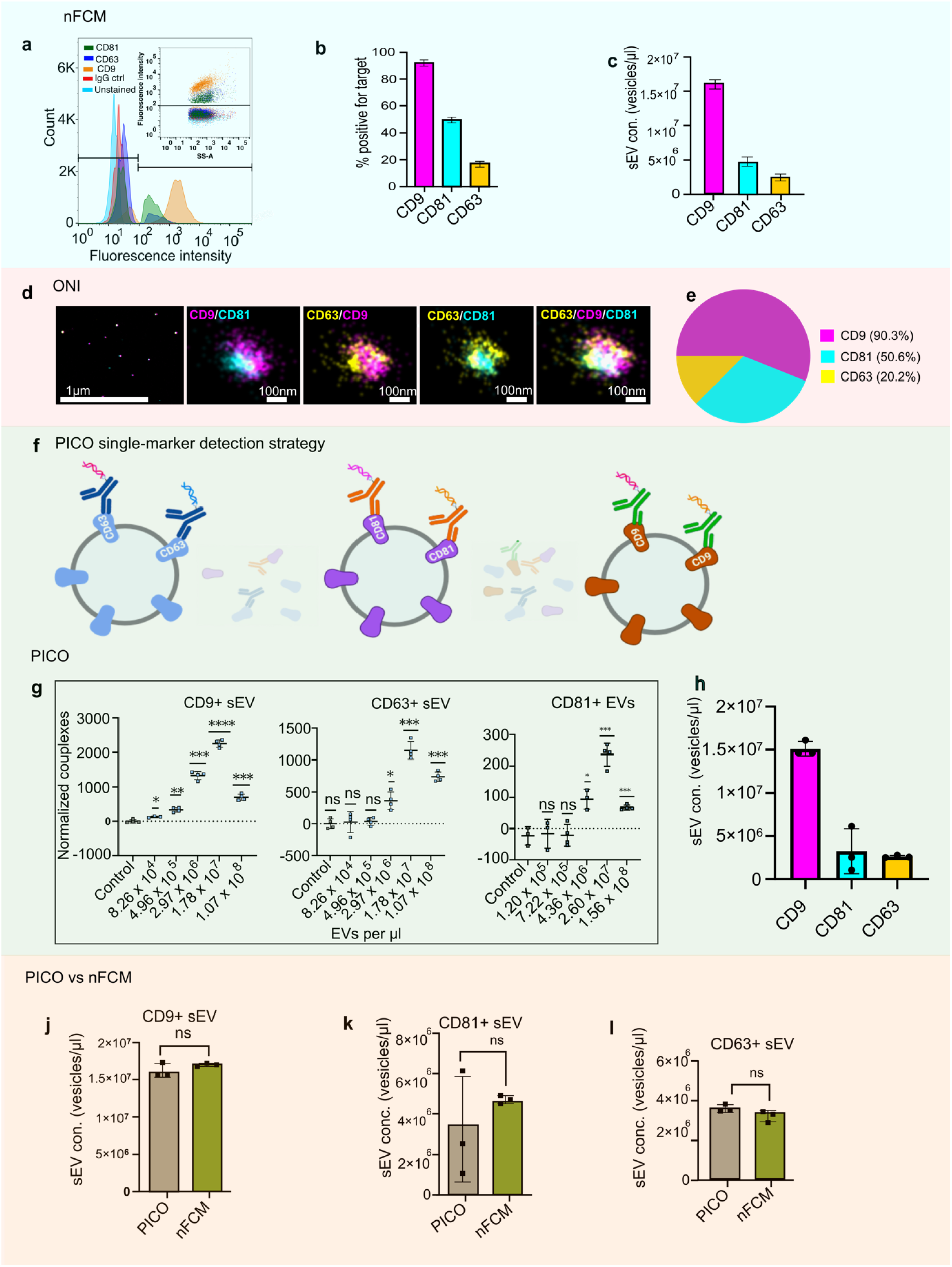
PICO enables absolute single-vesicle quantification of sEV positive for individual tetraspanins. ***a-c***, Single-vesicle analysis of HT1080-CD9 sEV by nanoflow cytometry (nFCM). Representative scatter plots **(a)** and quantification **(b)** of CD9, CD81, and CD63 positivity. Absolute concentrations of each subpopulation are shown in **(c)**. Data are presented as mean ± s.d. (n=3 biological replicates). **d,e,** Super-resolution dSTORM imaging validates tetraspanin expression. Representative dSTORM images of single sEV stained for CD9, CD81, and CD63 **(d)**. Quantification of the percentage of sEV positive for each marker from dSTORM clustering analysis **(e)**. **f,** Schematic of the PICO assay for detecting colocalized copies of the same protein on a single vesicle. Two monoclonal antibodies of the same clone, targeting a specific tetraspanin (e.g., CD9), are conjugated to distinct oligonucleotides. Following binding to a vesicle, sample partitioning and amplification generates a fluorescent couplex signal, indicating the presence of different copies of the target molecule on the same vesicle. **g,** Dilution curves for the single-marker PICO assay. Serial dilutions of HT1080-CD9 sEV incubated with antibody pairs (same clone) against CD9, CD81, or CD63. Corrected couplex counts, after normalization and subtraction of the antibody binding control (ABC), are plotted against sEV concentration. **h,** Absolute quantification of CD9⁺, CD81⁺, and CD63⁺ sEV subpopulations in HT1080-CD9 samples using the PICO assay. Data are presented as mean ± s.d. (n= 2 biological replicates). **j-l,** Correlation of absolute vesicle concentrations for CD9⁺ **(j)**, CD81⁺ **(k)**, and CD63⁺ **(l)** sEVs measured by PICO and nanoflow cytometry (nFCM).

### PICO enables single-vesicle marker co-localization to quantify sEV subpopulations

Clinical translation of EV biomarkers requires quantitative single-vesicle assays that can resolve cellular origin and quantify disease-associated EV subpopulations in complex samples; therefore, we evaluated the applicability of PICO for subpopulation analysis based on single-vesicle marker co-localization. As an orthogonal reference, we first characterized tetraspanin co-localization at single-vesicle resolution using nano-flow cytometry (nFCM). HT1080-CD9–derived sEV were dual-labeled with antibody pairs targeting CD9/CD81, CD9/CD63 or CD63/CD81. CD9^+^/CD81^+^ vesicles constituted the largest subpopulation (46.4%), followed by CD9^+^/CD63^+^ (19.7%) and CD63^+^/CD81^+^ (13.7%) (Fig. 4a-d), confirming the presence of distinct sEV subpopulations defined by marker co-expression. To enable quantitative subpopulation analysis based on single-vesicle marker co-localization, we adapted PICO to detect combinations of surface markers present on individual sEV. This was achieved by using antibody pairs as the fundamental detection unit and extending the approach to higher-order marker combinations (for example, CD9/CD63/CD81), with each antibody conjugated to a distinct oligonucleotide barcode (Fig. 4e). Because individual vesicles can bind multiple antibodies and each antibody species carries a unique oligonucleotide, partitions containing vesicles that present several surface markers generate fluorescent signals in multiple detection channels. Importantly, PICO operates under low partition occupancy (λ = 0.15 antibody molecules per partition), such that the probability of random co-localization of three or more distinct antibody species within the same partition is <1%, as predicted by Poisson statistics for digital PCR^28^. Accordingly, partitions exhibiting multiple fluorescent signals reflect genuine marker co-localization on individual vesicles rather than stochastic antibody co-partitioning, enabling robust quantitative delineation of EV subpopulations. Dilution curves were generated for each antibody pair across a range of sEV concentrations, with a peak *couplex* signal at 1.78 × 10⁷ sEV/µl (Fig. 4f-h). The relative abundances of sEV subpopulations detected by PICO mirrored those obtained by nFCM: CD9^+^/CD81^+^ events were most frequent, followed by CD9^+^/CD63^+^ and CD63^+^/CD81^+^. Triple-positive (CD9^+^/CD63^+^/CD81^+^) sEV were rare in both dSTORM and PICO analyses indicating that only a small fraction of vesicles carry all three markers (Fig. 4i, Fig. 3d). Absolute counts of marker-positive sEV showed no significant differences when compared to independent nFCM measurements of the same samples (Fig. 4j-l). As observed for single-marker detection, the coefficient of variation for marker co-localization measurements remained below 20% across technical and biological replicates (Extended Data Fig. 3a). Using the CD9/CD63 marker pair, the limit of detection (LOD) was 1.11 × 10⁶ sEV/µl and the limit of quantification (LOQ) was 3.68 × 10⁶ sEV/µl (Extended Data Fig. 3b). HT1080-derived sEV, which lack CD9 expression, showed no significant CD9/CD63 *couplex* signal relative to antibody-only controls, confirming assay specificity (Extended Data Fig. 3c).

**Figure 4.**
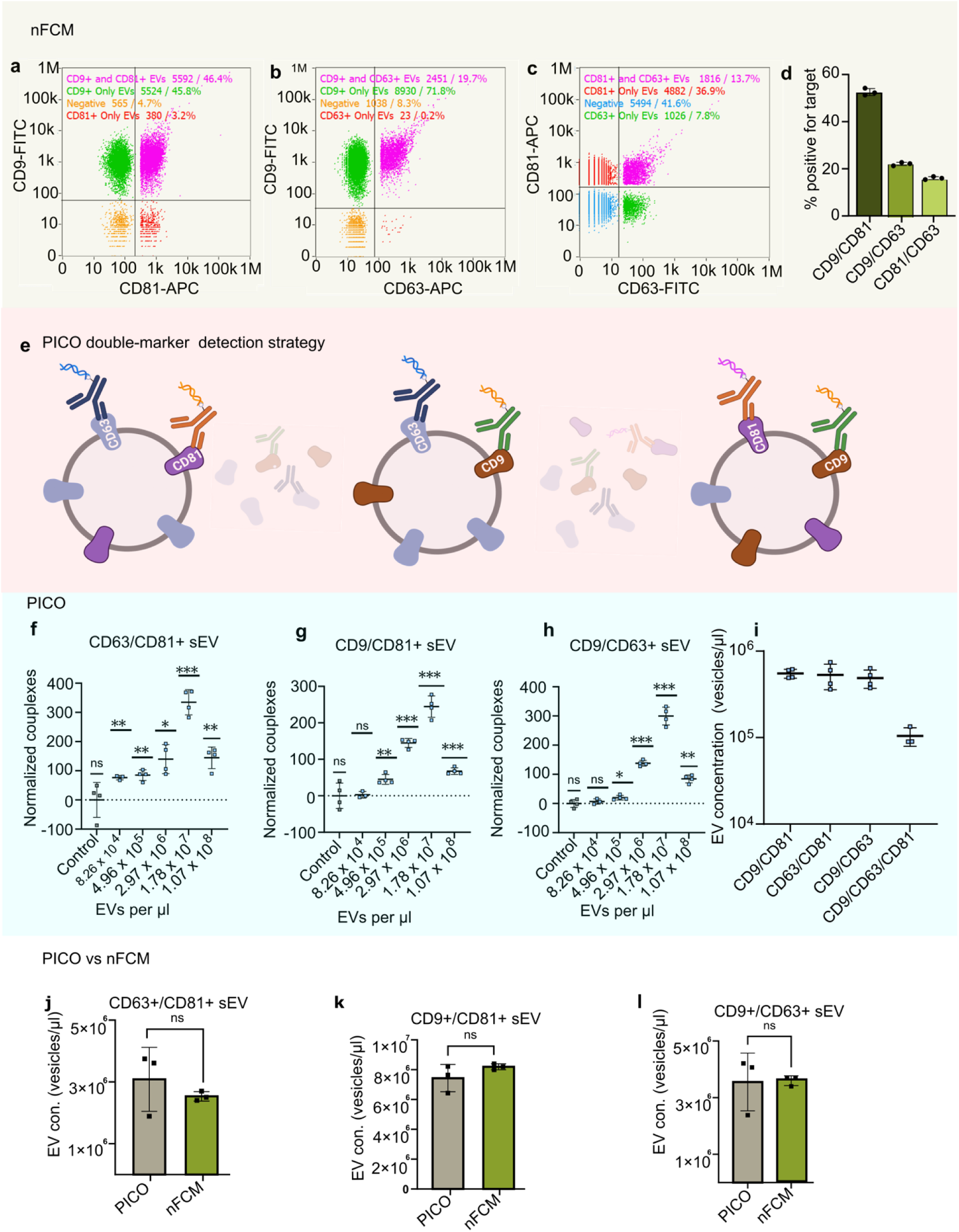
Single-vesicle marker co-localization enables absolute quantification of sEV subpopulations by PICO. ***a-d***, Single-vesicle analysis of HT1080-CD9 sEV by nanoflow cytometry (nFCM). Representative scatter plots for dual labeling with antibody pairs against CD9/CD81 (**a**), CD9/CD63 (**b**) and CD63/CD81 (**c**). The relative abundance of sEV subpopulations defined by co-localization of two markers is shown in (**d**). **e,** Schematic of the co-localization PICO assay. Two distinct antibodies (for example, anti-CD9 and anti-CD81), each conjugated to a unique oligonucleotide, bind their respective targets on the same vesicle. Sample partitioning and dPCR amplification generate a fluorescent readout corresponding to marker co-localization on single vesicles. **f-h,** Dilution curves for marker co-localization PICO detection. Serial dilutions of HT1080-CD9 sEV were incubated with antibody pairs against CD9/CD81 (**f**), CD9/CD63 (**g**) and CD63/CD81 (**h**). Corrected couplex counts, after normalization and subtraction of the antibody-only binding control (ABC), are plotted against sEV concentration. **i,** Quantification of sEV subpopulations defined by marker co-localization as detected by PICO; the inferred triple-positive (CD9^+^/CD63^+^/CD81^+^) population is also shown. J-l, Comparison of absolute concentrations of CD63⁺/CD81⁺ (j), CD9⁺/CD81⁺ (k) and CD9⁺/CD63⁺ (l) sEV subpopulations measured by PICO and nFCM. Data are presented as mean ± s.d. (n = 2 biological replicates); n.s., not significant (unpaired two-tailed t-test).

### Evaluation of sEV membrane integrity and detection of intravesicular protein markers using PICO

A prerequisite for single-vesicle analysis using PICO is that the assay conditions preserve the structural integrity of extracellular vesicles during antibody binding and processing. We therefore first assessed whether incubation of sEV with buffers used in the PICO workflow affects vesicle membrane integrity. HT1080-CD9–derived sEVs from the same preparation were incubated for 2 h either in PBS (reference condition), in PICO mastermix (containing all reagents used prior to sample partitioning), or in PICO lysis buffer, a detergent-containing formulation supplied for cell lysis and membrane protein extraction (Fig. 5a). Cryogenic electron microscopy (cryo-EM) revealed that sEV incubated in PBS or PICO mastermix retained intact lipid bilayer membranes (Fig. 5b, c). In sEV treated with PICO lysis the vesicular outline remained visible, whereas the lipid bilayer was no longer discernible, indicating loss of membrane integrity without complete collapse of vesicle morphology (Fig. 5d). To complement these structural observations and to test whether PICO can be configured to detect intravesicular proteins, we quantified a luminal sEV cargo protein under the same conditions. We selected eukaryotic translation initiation factor 4E-binding protein 1 (4E-BP1), a downstream effector of mTOR signaling reported to be present within sEVs, as a representative intravesicular marker^26^. Consistent with preserved vesicle integrity, 4E-BP1 was not detectable in intact sEV incubated in PBS or PICO mastermix (Fig. 5e). By contrast, robust detection and quantification of total 4E-BP1 were achieved following incubation in the PICO lysis buffer (Fig. 5f,g), confirming efficient release and accessibility of luminal cargo upon membrane disruption. Together, these results demonstrate that PICO preserves sEV membrane integrity under standard assay conditions for surface marker analysis, while enabling controlled detection of intravesicular proteins when vesicles are intentionally lysed. This modularity allows PICO to selectively interrogate either surface-exposed or luminal EV markers depending on the assay configuration.

**Figure 5.**
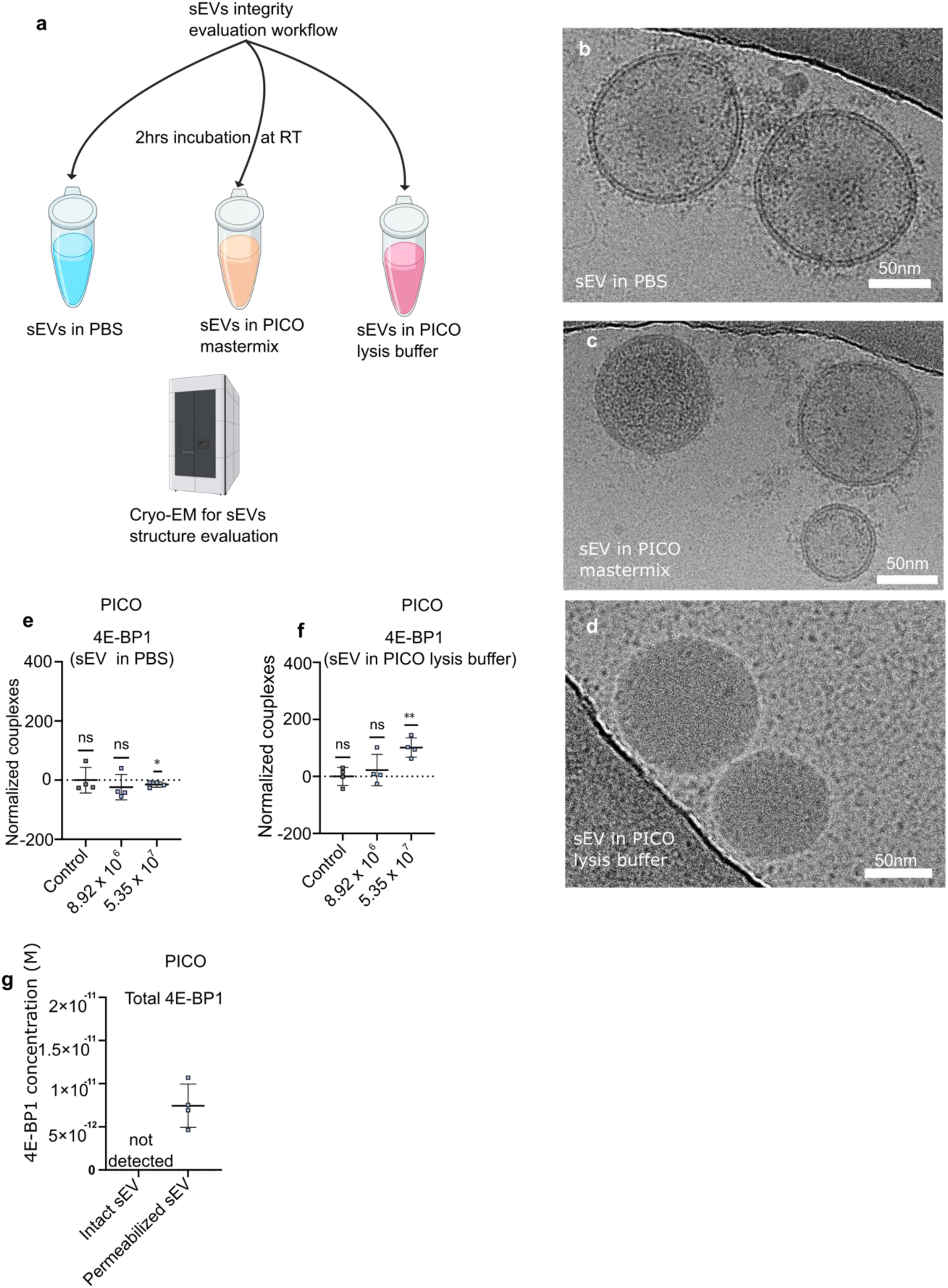
Buffer-controlled preservation and disruption of sEV integrity enables surface versus luminal readouts by PICO. **a**, Schematic of the experimental workflow. HT1080-CD9-derived sEV from the same batch were incubated for 2 hours in PBS (control), PICO mastermix (reaction buffer), or PICO lysis buffer (detergent-based). **b-d,** Representative cryo-electron microscopy (cryo-EM) images of sEV after incubation in PBS **(b)**, PICO mastermix **(c)**, or PICO lysis buffer **(d)**. Vesicles in PBS and mastermix retain intact lipid bilayers, while lysis buffer treatment results in loss of membrane integrity without complete collapse of vesicle morphology. Scale bars, 50nm. **e,f,** Quantification of the intravesicular protein 4E-BP1 by PICO. The target was undetectable in intact sEV incubated in PBS prior the reaction **(e)** but was robustly quantified after membrane permeabilization with PICO lysis buffer **(f)**. **g,** Absolute quantification of total 4E-BP1. Data are presented as mean ± s.d. (n=3 biological replicates).

### Quantitative co-localization defines cancer biomarker-positive sEV subpopulations by PICO

Moving beyond canonical tetraspanins, we next evaluated whether PICO can be applied to quantify oncology-relevant protein markers on sEV. This analysis requires subpopulation-resolved EV analysis, as clinically relevant signals often arise from specific subsets rather than from the total EV pool. This is particularly relevant in oncology, as HER2-targeted therapy has demonstrated clinical benefit even in tumors with low HER2 expression^29^. In parallel, we recently reported that full-length HER2 is detectable on a portion of sEV in breast cancer patient samples, supporting quantification of the HER2^+^ EV subpopulation as a promising route for liquid biopsy-based therapy monitoring^30^.

To test PICO for detection of HER2^+^ sEV, we used the HER2^+^ breast adenocarcinoma cell line MDA-MB-361. sEV were isolated from conditioned medium using the workflow in Fig. 2a and characterized by nFCM and cryo-EM. The preparation contained 6.52 × 10¹⁰ vesicles/mL with diameters of approximately 70 nm, and cryo-EM confirmed intact vesicular morphology with preserved lipid bilayers (Extended Data Fig. 4a,b). Immunoblotting detected CD9, CD63 and HER2 (Fig. 6b). Single-vesicle nFCM showed HER2 on 28.9% of sEVs (2.5 × 10¹⁰ vesicles/mL), with CD9 and CD63 on 59.7% (3.4 × 10¹⁰ vesicles/mL) and 25.8% (1.8 × 10¹⁰ vesicles/mL) of vesicles, respectively; HER3 served as a negative control (Fig. 6c,d; Extended Data Fig. 4c). We next applied PICO single-marker detection to define the vesicle concentration range supporting efficient *couplex* formation (Fig. 6e). PICO robustly detected CD9^+^, CD63^+^ and HER2^+^ sEV, consistent with immunoblotting and nFCM (Fig. 6b–d), whereas HER3 yielded no detectable signal, supporting assay specificity (Extended Data Fig. 4d). Quantification of HER2+ sEV by PICO yielded 5.0 × 10⁶ vesicles/µl, closely matching nFCM measurements (6.0 × 10⁶ vesicles/µl) (Fig. 6f). To resolve HER2-associated sEV subpopulations, we quantified marker co-localization with canonical tetraspanins by nFCM. HER2 was confined to a subset of vesicles and showed a strongly non-uniform distribution across tetraspanin-defined fractions: HER2⁺CD9⁺ vesicles accounted for 18.6% (3.2 × 10⁶ vesicles/µl), whereas HER2⁺CD63⁺ vesicles were rare (0.3%; 1.4 × 10⁴ vesicles/µl); CD9⁺CD63⁺ vesicles represented 10.6% (1.9 × 10⁶ vesicles/µl) (Fig. 6g–i). These co-localization patterns underscore the intrinsic heterogeneity of sEV populations and illustrate that clinically relevant biomarkers may reside within specific vesicle subsets rather than the bulk EV pool. We then implemented PICO co-localization assays using antibody pairs targeting CD9/CD63 and HER2/CD9, with HER2/HER3 and HER3/CD9 included as specificity controls. Dilution series demonstrated consistent performance across sEV concentrations (Fig. 6j), with *couplex* signals detectable down to 10⁵ sEVs/µl, in line with sensitivity benchmarks established using HT1080-derived reference sEV; HER3-containing pairs yielded no significant *couplex* signal (Extended Data Fig. 4e,f). Quantification of HER2⁺CD9⁺ sEV by PICO closely matched nFCM measurements (Fig. 6k), supporting scalable, subpopulation-resolved EV readouts as a key requirement for diagnostically robust liquid biopsy assays.

**Figure 6.**
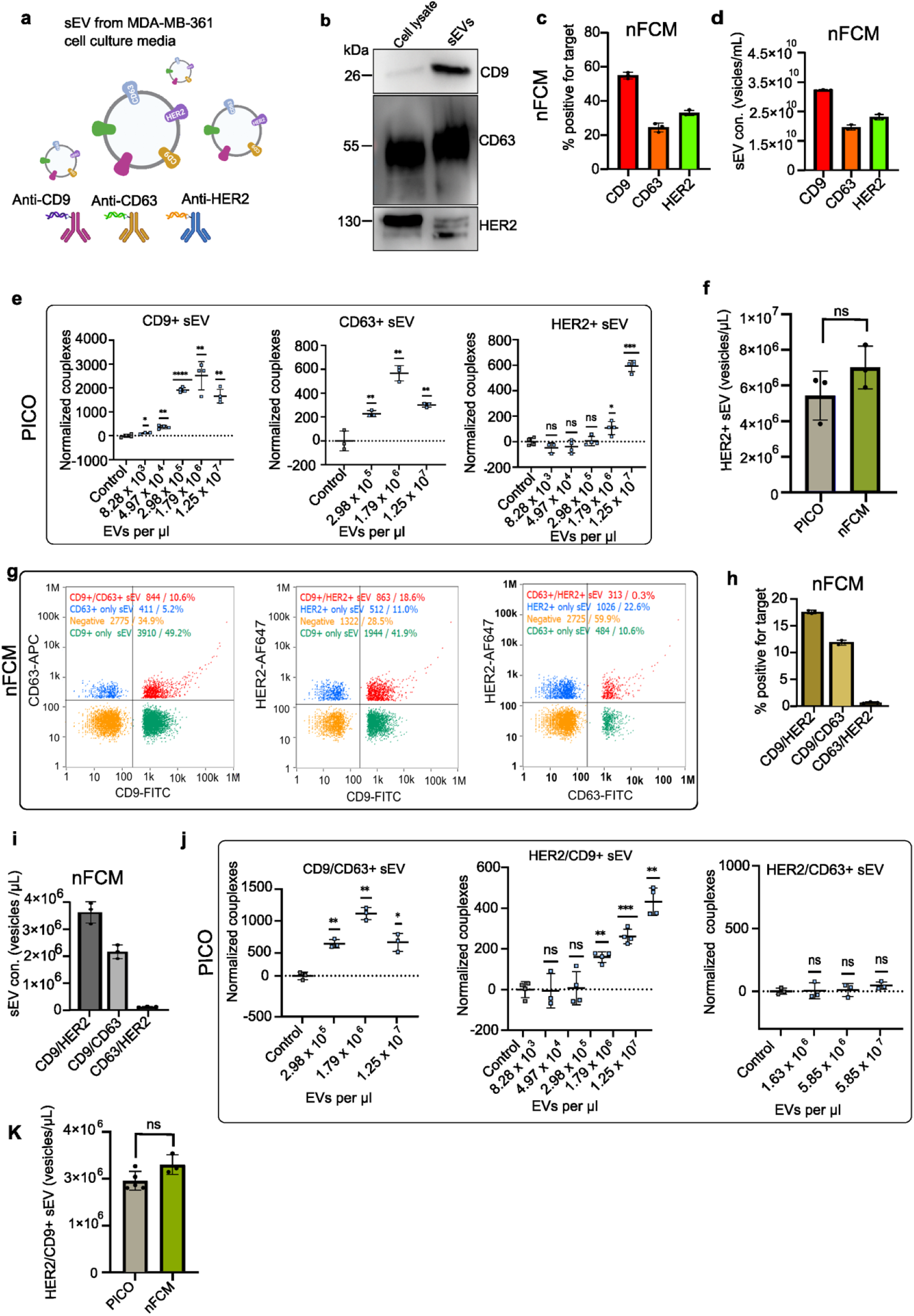
sEV subpopulation analysis by PICO using quantitative colocalization analysis. **a**, Schematic of sEV derived from the HER2^+^ brain metastasis-derived breast cancer cell line MDA-MB-361 **b,** immunoblot analysis of MDA-MB-361-derived sEV probed for CD9, CD63, HER2. **c, d,** Single-vesicle analysis of MDA-MB-361-derived sEV by nFCM. **c,** Quantification of the percentage CD9^+^, CD63^+^, HER2+ and **d,** concentration of sEVs positive for each marker. **e,** PICO single-marker dilution series for CD9, CD63, and HER2 to determine the optimal sEV concentration for efficient couplex formation. **f,** Quantification of HER2^+^ sEV concentration by PICO and nFCM. **g-i,** Analysis of biomarker co-expression on sEV using PICO. Representative images (**g**) and quantification of sEV subpopulations co-expressing CD9/CD63 (**i**), HER2/CD9 (**j**), and HER2/CD63 (**k**). **l-n,** Dilution series of PICO for detection of marker pairs CD9/CD63 (**l**), HER2/CD9 (**m**), and HER2/CD63 (**n**). **n,** Comparison of couplex counts for different marker pairs (CD9/HER2, CD9/CD63, CD63/HER2) measured by PICO, with a trend corroborated by nFCM. **o,** Quantification of HER2^+^/CD9^+^ sEV by PICO and nFCM, showing no significant difference (two-tailed unpaired t-test; ns, not significant). All nFCM (n=3) and PICO(n=2) (data in panels d, g, i-k, o, and p are from biological replicates and are presented as mean ± s.d.

### PICO discriminates breast cancer patient–derived sEV from healthy donors by quantitative co-localization analysis

We next assessed the clinical applicability of PICO for subpopulation-resolved diagnostics using plasma from patients with HER2-positive breast cancer (n = 4) and matched healthy donors (n = 4). sEVs were isolated by OptiPrep density gradient centrifugation (nine fractions; Fig. 7a); EV-enriched fractions 2–3 were identified by NTA and immunoblotting (particle counts and CD9/HER2 signal) and pooled for downstream analyses (Fig. 7b). Breast cancer samples showed a modestly higher vesicle concentration than controls (5.6 × 10¹⁰ vs 4.0 × 10¹⁰ vesicles/mL), with comparable size distributions (Fig. 7c; Extended Data Fig. 5a), and cryo-EM confirmed intact bilayer morphology in all samples (Fig. 7d). Single-vesicle nFCM revealed a higher proportion of CD9⁺ sEVs in breast cancer–derived samples (51%; 2.5 × 10⁶ vesicles/µL) than in healthy donors (15%; 1.3 × 10⁶ vesicles/µL), whereas CD63⁺ fractions were comparable between cohorts. Notably, HER2⁺ sEVs were detected only in breast cancer samples (15%; 1.4 × 10⁶ vesicles/µL) and were not observed in healthy donors. HER3 remained undetectable in both groups (Fig. 7e; Extended Data Fig. 5b). We then applied PICO single-marker detection to the same samples. Dilution series were performed to define optimal sEV input concentrations for absolute quantification. Consistent with nFCM, CD9⁺ and CD63⁺ sEV were detected in both cohorts, whereas HER2⁺ sEV were observed exclusively in breast cancer samples. HER3 yielded no detectable *couplex* signals in any sample (Fig. 7f; Extended Data Fig. 5c). Absolute quantification by PICO closely matched nFCM measurements across all tested markers (Fig. 7g). To assess diagnostically relevant subpopulations, we next analyzed marker co-localization. nFCM revealed substantial enrichment of HER2⁺CD9⁺ (11% vs. 0.5%) and HER2⁺CD63⁺ (4.4% vs. 0.8%) sEV in cancer samples compared to healthy donors, whereas CD9⁺CD63⁺ co-expression was similar (9% vs. 7%) (Fig. 7h; Extended Data Fig. 6a). Absolute counts corresponded to 1.4 × 10⁶ (HER2⁺CD9⁺) and 4.1 × 10⁵ (HER2⁺CD63⁺) vesicles/µL in breast cancer samples, with no detectable HER2-associated subpopulations in healthy donors. We then performed PICO co-localization analysis to quantify defined sEV subpopulations (CD9⁺CD63⁺, HER2⁺CD9⁺ and HER2⁺CD63⁺). Consistent with nFCM, all three subpopulations were detected in breast cancer samples, whereas healthy donors displayed only CD9⁺CD63⁺ vesicles, reflecting the absence of HER2 (Fig. 7i; Extended Data Fig. 6b). Lack of HER2 binding eliminated *couplex* formation in control samples, confirming assay specificity. PICO-derived absolute counts were concordant with nFCM measurements (Fig. 7j). Collectively, these data demonstrate that quantitative co-localization analysis by PICO robustly discriminates breast cancer-derived sEV from healthy donor samples by identifying HER2-defined vesicle subpopulations. Importantly, this approach resolves tumor-associated EV subsets against a heterogeneous background of circulating vesicles, highlighting its potential for clinically actionable liquid biopsy diagnostics.

**Figure 7.**
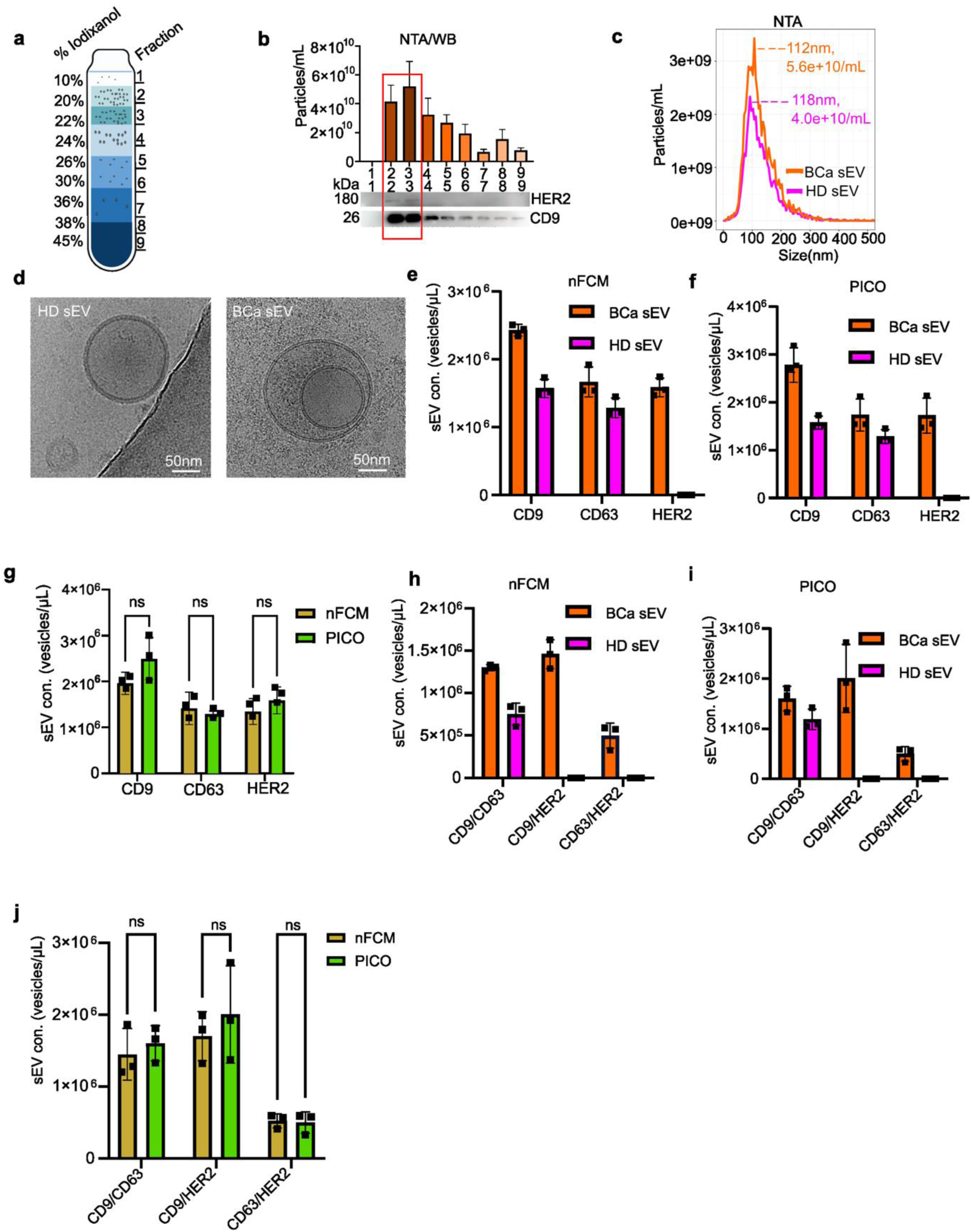
Single-vesicle PICO analysis distinguishes breast cancer patients from healthy donors by quantitative marker co-localization. **a**, Schematic of plasma sEV isolation from breast cancer patients (n = 4) and matched healthy donors (n = 4) using an OptiPrep density gradient. Nine fractions were collected. **b**, Characterization of gradient fractions. NTA particle concentration (left y-axis, bars) and immunoblotting (lower panels) for CD9 and HER2 across fractions. Fractions 2–3, enriched in sEV as indicated by high particle counts and CD9/HER2 signal, were pooled for downstream analyses. **c**, Representative NTA size distribution profiles of purified plasma sEV from a breast cancer patient (BCa) and a matched healthy donor (HD). **d**, Representative cryo-EM images of purified plasma sEV from a breast cancer patient and a healthy donor, showing intact vesicular morphology with preserved lipid bilayers. Scale bars, 50 nm. **e**, nFCM-based single-vesicle quantification of marker-defined sEV subpopulations. Bars show the mean concentration (vesicles/µL) of CD9⁺, CD63⁺ or HER2⁺ sEV in breast cancer and healthy donor samples. HER3 served as a negative control (Extended Data Fig. 5b). Data are mean ± s.d.; n = 4 biologically independent samples per group. **f,** PICO single-marker analysis of the same sEV preparations as in e, quantifying CD9⁺, CD63⁺ and HER2⁺ sEV subpopulations. HER3 was not detected (Extended Data Fig. 5c). **g**, Head-to-head comparison of absolute sEV counts for CD9⁺, CD63⁺ and HER2⁺ subpopulations measured by nFCM and PICO. Data are mean ± s.d.; n = 2 biologically independent samples per group; two-tailed unpaired t-test, n.s. **h**, nFCM-based quantitative co-localization analysis of sEV subpopulations defined by marker combinations (CD9⁺CD63⁺, HER2⁺CD9⁺, HER2⁺CD63⁺). Data are mean ± s.d.; n = 4 biologically independent samples per group. **i,** PICO co-localization analysis of the same marker-defined sEV subpopulations as in h (CD9⁺CD63⁺, HER2⁺CD9⁺, HER2⁺CD63⁺). No HER2-associated subpopulation was detected in healthy donor samples. **j**, Head-to-head comparison of absolute sEV counts for co-localization defined subpopulations measured by nFCM and PICO. Data are mean ± s.d.; n = 2 biologically independent samples per group; two-tailed unpaired t-test, n.s

## Discussion

Progress toward reliable EV-based diagnostics has been limited by the absence of methods that provide absolute quantification and high specificity while enabling robust biomarker profiling at the single-vesicle level in complex biofluids in a format that is simple to implement and compatible with high-throughput workflows ^31,30,32,33^. Here, we present a new application of PICO, a digital immunoassay that addresses these challenges by enabling reference-free measurements with single-vesicle resolution for surface markers and quantitative detection of intravesicular cargo. The core innovation of PICO is a dual-labeling design in which a detectable signal is generated only when two independent antibody labels co-localize on the same physical entity. This co-localization requirement strongly suppresses background from unbound reagents and soluble species, thereby reducing a major source of false-positive events that can complicate single-vesicle flow cytometry and conventional immunoassays such as ELISA^34^. While large, non-vesicular aggregates bearing multiple copies of a target could, in principle, generate a signal, our experimental controls indicate that the assay is not measurably confounded by free soluble antigens, which represent the dominant background in biological samples. The requirement for dual-antibody co-localization favors multivalent membrane-bound structures and thereby enhances specificity for vesicle-associated targets. As a result, the digital *couplex* readout functions as a highly specific reporter of vesicle-bound biomarkers rather than soluble species, which is, to our knowledge, largely unique feature of the PICO assay design. We further benchmarked PICO against established orthogonal technologies to position it within the current analytical landscape. The strong concordance with nanoflow cytometry (nFCM) across multiple markers and sample types supports the quantitative accuracy of PICO for absolute subpopulation measurements. At the same time, PICO provides distinct advantages in sensitivity and multiplexing. The assay achieved a limit of detection of approximately (∼1.15 × 10^6^ sEV/µL) for abundant biomarker-defined subpopulations, reaching biomarker-specific sensitivity comparable to fluorescence-based nFCM, which typically operates in the range of 10⁶–10⁷ particles/mL. However, fluorescence-based nFCM measurements can be affected by background signal, including autofluorescence and the limited effective brightness of fluorophores on nanoscale particles, which together can obscure weakly labeled subpopulations ^22,23^ In addition, detection of low-abundance biomarkers is constrained by the need for multiple target copies per vesicle to exceed the fluorescence detection threshold; published estimates suggest that on the order of ∼9 or more copies may be required under typical conditions^35–37^. Similar limitations of fluorescence-based detection on nanoscale particles have also been reported for fluorescent NTA (fNTA)^38^. By contrast, PICO can register vesicle-associated targets at substantially lower copy numbers—down to a single copy per vesicle in the *couplex* format (and two copies for single-marker detection,). PICO’s DNA-barcoded readout, decoded by dPCR, supports multiplexed single-vesicle analysis up to the instrument’s channel capacity, enabling quantification of co-localization-defined subpopulations, such as CD9/CD63/CD81 co-expression patterns and corresponding double-positive vesicle fractions demonstrated in this work. Application of PICO to well-characterized reference materials revealed an increase in CD9⁺ vesicle populations that was consistent with measurements obtained by NTA and nFCM, thereby validating across orthogonal platforms the observation that CD9 expression promotes EV production, as described for several tetraspanins. ^36,39,40^. Furthermore, orthogonal single-vesicle analyses, including dSTORM, underscore the heterogeneity of sEV populations, with substantial vesicle-to-vesicle variation in marker composition and cargo. This is consistent with a growing body of work from emerging high-resolution modalities, including single-particle proteomics and imaging mass cytometry^41^. Importantly, PICO turns an intrinsic property of Evs - the multivalent presentation of epitopes on a membrane surface - into an analytical advantage: by requiring co-localization of two antibody labels on the same physical structure, it enables digital counting of vesicle-bound targets without dependence on fluorescence intensity scaling, equilibrium-based signal fitting, or external calibration curves. A significant and distinctive capability of PICO is its tunable access to the intravesicular lumen. Under standard assay conditions, sEV membrane integrity is maintained, as supported by cryo-EM and by the absence of detectable luminal 4E-BP1. Upon controlled permeabilization, the same workflow enables robust quantification of intravesicular cargo. This integrated assessment of both membrane and luminal markers within a single quantitative platform extends beyond methods that are largely confined to surface phenotyping. Moreover, intravesicular cargo measurements provide an intrinsic quality-control readout: luminal contents are released upon vesicle disruption (for example, after repeated freeze–thaw cycles), such that reduced cargo signal can serve as an indirect indicator of sample integrity. We next applied PICO to detect and quantify the tumor-associated marker HER2 on sEV derived from cell lines and from patient plasma. Diagnostic discrimination improved when analysis focused on marker-defined subpopulations rather than single-marker positivity alone. A similar pattern was observed in the cell-model reference system, where HER2 preferentially co-localized with CD9, while the CD63⁺ subset - despite comprising 25.8% of the total sEV population - contained no detectable HER2. In patient plasma, HER2/CD9 double-positive vesicles were more abundant than HER2/CD63 double-positive vesicles (11% versus 4.4%). In contrast, the CD9/CD63 double-positive population was comparable between patients and healthy donors, indicating that disease specificity arises from tumor-marker defined subpopulations rather than from general tetraspanin co-expression. This moves beyond merely detecting a cancer-associated marker to identifying specific, disease-relevant vesicle populations, a crucial step for developing robust liquid biopsy tests that can stratify patients^42^. The strong correlation between PICO and nFCM across all tested clinical samples underscores the technical robustness and immediate utility of the platform for analyzing complex, low-abundance clinical specimens. Despite its strengths, certain considerations of the PICO platform warrant discussion. Several limitations of the current PICO implementation warrant consideration. As with all dual-antibody immunoassays, PICO is susceptible to the Hook effect (antigen excess phenomenon). Under increasing target concentrations, *couplex* formation initially rises but declines at very high analyte levels, where antibody saturation prevents simultaneous binding of two antibodies to the same target. This non-linear behavior is observable in sEV measurements (e.g., Fig. 3f,g). Although mathematical correction frameworks for antigen-excess kinetics have been proposed, such models have not yet been adapted for PICO-based sEV quantification. Accordingly, quantitative analyses in this study were restricted to the linear (“low-side”) regime of the response curve to avoid underestimation. Multiplexing capacity is currently limited by the number of available dPCR instrument detection channels, although future implementations could expand this through combinatorial barcoding or amplitude-based strategies. As with any antibody-dependent platform, assay performance is contingent on the availability of high-quality monoclonal antibodies, which may limit implementation for poorly characterized targets. Finally, while detection of triple-positive vesicles is technically feasible, its sensitivity is constrained by partition number and target abundance. In the present study, triple-positive sEV concentrations approached the lower detection range (∼1.04 × 10⁵ sEV/µL), where reliable discrimination becomes challenging. Increasing dPCR partition density would enhance sensitivity for higher-order multiplexing. Addressing these challenges represents an important direction for future development of the PICO platform; however, systematic optimization in this regard was beyond the scope of the present study. In summary, we present a new application of PICO that enables reference-free, single-vesicle profiling of extracellular vesicles in complex biofluids. By coupling biomarker co-localization on individual vesicles to a ddPCR-based digital readout, PICO provides absolute, multiplexed quantification without reliance on specialized optical instrumentation or calibration workflows. This format brings single-vesicle EV analysis closer to routine laboratory implementation while retaining the sensitivity of digital PCR. We anticipate that PICO’s capacity to resolve EV heterogeneity and to quantify both surface and luminal markers will advance fundamental vesicle biology, accelerate biomarker discovery, and support the development of next-generation EV-based diagnostics.

## Methods

### Ethical Statement

Plasma samples used in this study were previously collected at the Department of Obstetrics and Gynecology, University of Freiburg Medical Center. All procedures involving human subjects were conducted in compliance with international ethical guidelines and were approved by the University of Freiburg’s Institutional Review Board (Protocol No. 36/12). Written informed consent was obtained from all participants or their legal representatives prior to sample collection. To ensure privacy and confidentiality, all samples provided to our research team were fully deidentified.

### Cell culture

HT1080 (CCL-121) and MDA-MB-361(HTB-27) (purchased from American Type Cell Collection) were maintained under standard mammalian cell culture conditions (37°C, 5% CO₂, humidified atmosphere). HT1080 cells were grown in RPMI 1640 (1X) medium (Gibco, 21875-034) supplemented with 10% heat-inactivated fetal bovine serum (FBS; PAN BIOTECH, P30-1506). HT1080-CD9 were maintained with RPMI 1640 (1X) medium (Gibco, 21875-034) supplemented with 10% heat-inactivated fetal bovine serum (FBS; PAN BIOTECH, P30-1506) and 200 µg/mL hygromycin B (Invitrogen, 10687010). MDA-MB-361 cells were maintained with DMEM (1X) (Gibco, 41966029) supplemented with 10% FBS.

### Generation of HT1080-CD9 stable cell line

HT1080 cells were trypsinized from a confluent T-75 flask and counted. 4 x 10^4^ cells/ well were seeded in a 24-well plate in 500μl complete growth medium. On the next day, cell confluency was controlled. 70-80% confluent cells were chosen for carrying out transfection. For transfection Lipofectamine-LTX was used. The transfection procedure was carried out according to the recommendations of the supplier. For each well of cells to be transfected, 0.5μg of plasmid DNA (PCDNA3, Invitrogen, V87020) was diluted into 50μl of Opti-MEM reduced serum medium and mixed gently. The plasmid DNA dilution was carried out in a 96-well plate. PLUS, reagent was mixed gently before use and then 0.5μl of which was added (a 1:1 ratio to DNA) directly to the diluted DNA. The mixture was mixed gently and incubated for 5 min at room temperature. Lipofectamine-LTX transfection reagent was mixed gently before use. For each well of cells, 1.25μl of Lipofectamine-LTX was diluted in 50μl of Opti-MEM reduced serum medium, mixed gently and then added to the above diluted DNA solution, mixed gently and incubated for 25 min at room temperature. For generation of stable cell lines, complete growth medium, supplemented with Hygromycin, 200μg/mL was added on the transfected cells the day after transfection. The cells were kept under selection, with medium change every three days, for about 7-10 days. Cells expressing high level of CD9 were sorted in core facility-Universitätklinikum-Freiburg with using (Becton Dickinson),

### Flow cytometry analysis

The HT1080 and HT1080-CD9 cells were trypsinised and harvested with Trypsin-EDTA for a single cell suspension and were counted using a cytometer (Company, product number). For surface marker staining, cells were incubated for 1 hr at 37°C so as to revive all surface proteins. Cells were centrifuged at 1500 xg for 3 min and the supernatant was discarded. Cells were washed with cold PBS/1% BSA, centrifuged and supernatant discarded. Followed by this, 2.0×10^4^ cells were stained, respectively, with following fluorophore-labelled antibodies at 4°C for 25 min: CD9(-APC (H19a, Biolegend), CD81-APC (TAPA-1, Biolegend), CD63-APC (H5C6, Biolegend). Next, unbound antibodies were removed from the cells by performing a two-step wash with PBS, after which the cells were resuspended in PBS for further analysis using a flow cytometer (BD Accuri, product number). Data were processed using FlowJo software (version 10.10.0)

### Preparation of conditioned medium and sEV isolation

HT1080, HT1080-CD9 and MDA-MB-361 cells were seeded in 20x 14.5 cm culture dishes with complete medium. After 48 hours, 20 mL of conditioned cell culture medium from 20 x14.5 cm cell culture dish plates were collected in 50 mL falcons (400 mL), centrifuged at 8,000 x g for 5 min at 4 °C and then at 2,500 x g for 25 min at 4 °C. The supernatant was then filtered through 0.2 μm filter, to remove microvesicles and remaining cell debris. The resulting filtrate, containing soluble proteins and sEV, was concentrated by tangential flow filtration (TFF)-based devices (Hansabiomed, HBM-TFF-Easy), connected to a peristatic pump (Repligen, product numer) at 50 mL/min. The volume of the concentrated conditioned media after TFF was ∼1000 µL. The concentrated medium was collected into 1.5-mL tubes (Eppendorf, 0030108116). The concentrated conditioned media was divided into two volumes of 500µL for size exclusion chromatography (SEC). The EVs were further purified by SEC using 35-nm qEV columns (IZON, ICO-35) and an automatic fraction collector V2 (IZON, AFC-V2), according to the manufacturer’s instructions. Both the SEC columns and samples were brought to room temperature prior to use. The buffer volume was 2.91 mL, and the purified collection volume was 0.5 mL in one fraction on AFC-V2. The samples were loaded on the loading frit of SEC columns, and the buffer volume was collected immediately. The PBS filtered through 0.22-µm filters was topped-up when all the samples entered the frit. After the buffer volume reached 2.91 mL, the 0.5-mL EV samples were collected into 1.5-mL tubes (Eppendorf, 0030108132). A total of 10 fractions were collected. Purified EV samples were aliquoted and stored at −80°C if the samples were not used immediately.

### Immunoblotting

Cells or sEV were lysed using RIPA buffer (Thermo Fisher, 89900) and total protein was quantified using a microBCA protein quantification kit (Thermo Scientific, 23235). Subsequently, 30 μg andl 5 μg of cell lysate and sEV lysate protein respectively was separated by sodium dodecyl sulfate-polyacrylamide (12%) gel electrophoresis, transferred onto a PVDF membrane (Millipore, ISEQ85R). The membranes were blocked in 5% milk for one hour at room temperature, followed by 5 minutes three times washes with TBST. The PVDF membranes were then incubated overnight at 4◦C with primary antibodies against CD9 (Cell signaling Technology, D801A, 1:1000 dilution in 3% milk), CD63 (Cell Signaling Technology, E1W3T, 1:1000 dilution in 3% milk), CD81 (Cell Signaling Technology, D3N2D, 1:1000 dilution in 3% milk), Vinculin (Cell Signaling Technology, E1E9V, 1:1000 dilution in 3% milk), HER2 (Cell Signaling Technology, 29D8), GAPDH (Santa Cruz Technology, Sc-47724, 1:5000). Following 3X wahing steps with TBST, the PVDF membrane was incubated with the corresponding horseradish peroxidase-conjugated secondary antibody (Jackson Immuno Research, 115035003 and 111035003) at room temperature for one hour. Following another round of washing with TBST, protein bands were visualised with the FUSION FX7 imaging system (Vilber, fusion FX7).

### Nanoparticle Tracking Analysis (NTA)

Size distribution, concentration of sEVs were analyzed using a QUATT Nanoparticle Tracking Analyzer (Particle Metrix GmbH) equipped with a 488 nm laser. Measurements were performed in 0.1x PBS (prepared in Milli-Q H_2_O from an OmniaTap xs basic system, Stakpure). For size and concentration measurements in scatter mode, samples were diluted to achieve a particle count of 80–200 particles per frame. Settings were as follows: sensitivity 80, shutter 90, trace length 12s, frame rate 30 fps, 7 positions per sample, minimum expected particle size 10 nm, maximum expected particle size 1,000 nm, temperature 25 °C. Data were acquired using ZetaView Navigator software (v.1.4.2.1) and final size distribution histograms were generated using Phonups software.

### Preparation of cryo-EM samples and TEM Imaging

Holey carbon grids (Quantifoil R 1.2/1.3 Cu 300 mesh) were glow discharged for 25 seconds at 10 mA (Pelco EasiGlow). 3 ul of the purified sEV samples were pipetted onto the grid, blotted (Vitrobot Mark IV, 22°C, 100% humidity, 6 sec, force 5) and plunged frozen in liquid ethane. To increase the particle density on the grid when needed, multiple rounds of sample application and blotting were performed before the sample was plunged frozen. Samples were imaged on a 300 kV Krios G4 with Selectris Energy Filter and Falcon 4i direct electron detector. Standard parameters were: 105kx magnification, total dose 40 e^-^/Å², 3 µm defocus.

### DNA labeling of antibodies and PICO assay

The antibodies employed in this study were labeled in a specific site of the heavy chain with amplifiable DNA oligonucleotide labels using the PICO BioScience Antibody Labeling Service. To have a low limit of detection, an antibody concentration in the binding reaction of 4 x 10^-11^ M was employed, while for absolute quantification of EVs and proteins the antibody concentration in the binding reaction was increased to 5 x 10^-10^ M, as previously described^24^. For the detection of surface markers in intact EVs, the samples and antibodies were mixed in Control Buffer (PICO BioScience), while for intravesicular cargo in lysed EVs, the samples were first lysed in Lysis Buffer (PICO BioScience) for 3 h. In both cases, samples and antibodies were mixed (1μl each) overnight at 4 °C following the PICO Amplification Core Kit (#PICO-000010, PICO BioScience) instructions. In every experiment, an Antibody Binding Control (ABC) was included, which contains the same antibody mix in the appropriate buffer, but lacks the sample of interest. The signal detected in this control will be employed later during data processing. As the PICO assay always requires the use of at least two antibodies with different DNA labels, for the detection of EVs positive for an individual surface marker, a single monoclonal antibody was selected, divided in two pools, and each of them labeled with a different PICO Label. Before the dPCR amplification step, the samples were pre-diluted to aim for an average lambda of 0.15, as recommended by^24^. The dPCR was performed using QIAGEN’s QIAcuity Digital PCR System according to PICO BioScience’s manual, using the PICO Probes matching the DNA labels attached to the antibodies (PICO BL, P8, N6, and O7 probes; #PICO-000070 to 73, PICO BioScience). After priming, leading to the partitioning of the sample, 40 cycles are applied with a denaturing step at 95°C for 15 s and an annealing step at 58°C for 30 s. For imaging, for the green channel an exposure time of 500 ms and a gain of 6 was used, for the yellow channel an exposure time of 400 ms and a gain of 6 was used, for the orange channel an exposure time of 400 ms and a gain of 6 was used, and an exposure time of 300 ms and a gain of 4 was used for detection in the red channel. Each probe detects its corresponding label and generates a fluorophore detected in the associated channel: P8 label/probe lead to FAM signal (𝝀_ex_ = 493 nm, 𝝀_em_ = 517 nm) measured in the green channel, BL label/probe to HEX signal (𝝀_ex_ = 533 nm, 𝝀_em_ = 559 nm) measured in the yellow channel, N6 label/probe to Atto550 signal (𝝀_ex_ = 553 nm, 𝝀_em_ = 575 nm) in the orange channel, and O7 label/probe to Texas Red (𝝀_ex_ = 586 nm, 𝝀_em_ = 603 nm) measured in the red channel. It is important to note that the channel may vary depending on the instrument and the excitation wavelengths and detection filters used. The raw dPCR data was analyzed using PICO BioScience’s PI-QUANT software (https://pico-bioscience.shinyapps.io/piquant/). The raw couplexes were processed as described in^24^, incorporating both ABC correction and lambda normalization. ABC correction accounts for any offsets in the dPCR data, such as signal dropouts or incorrect clustering, while lambda normalization corrects for handling errors, assuming that technical replicates should share the same lambda value. When using saturation conditions, it can be assumed that the couplexes represent the EVs. Therefore, in order to calculate the molar concentration, the couplexes were multiplied with the dilution factor with which the sample was initially diluted for dPCR and the count of couplexes/EVs transferred in the unit of interest (e.g. copies/µl). This was used as absolute data. Applying the appropriate statistical test, it was assumed that the data followed a normal distribution based on the theoretical statistical distribution of couplexes.

### Nano-flow cytometry analysis

Nanoflow cytometry (nFCM) analysis was performed on a Nano Analyzer (NanoFCM Co., Ltd., Nottingham, UK) equipped with a 488 nm laser. The instrument was calibrated prior to measurements using 250 nm silica calibration beads (QS3003, NanoFCM Co.) at a concentration of 2.11 × 10¹⁰ particles/mL, which also served as a reference for determining particle concentration. Size calibration was performed using monodisperse silica beads (S16M-Exo, NanoFCM Co. Ltd.) of four different sizes (68 nm, 91 nm, 113 nm, 155 nm). Freshly filtered (0.22 µm) 1× PBS was analyzed as a background control, and its signal was subtracted from all subsequent sample measurements. Data were acquired over a 1-minute collection time at a constant sample pressure of 1.0 kPa. For analysis, sEVs samples were diluted in 0.1 µm-filtered 1× PBS to achieve a particle count within the optimal range of 2,500–12,000 events per minute. Particle concentration and size distribution were determined using the NF Profession v2.08 software (NanoFCM Co. Ltd.). For immunofluorescence staining, surface protein markers on sEVs (CD9, CD81, CD63, HER2, HER3). sEV samples were incubated with a cocktail of fluorescently conjugated antibodies, including Anti-human-CD9-APC (clone HI9a, 312108, Biolegend), Anti-human-CD63-APC (clone H5C6, 353007, Biolegend), Anti-human-CD81-APC (clone 5A6, 349509, Biolegend) and Anti-human-HER2-AF647 (clone 24D2, 616153, Biolegend), for 1 h at room temperature in the dark. Following incubation, samples were diluted in PBS for immediate analysis. To ensure signal specificity, the following controls were included blank control, antibody-only, detergent-lysed sEVs and isotype control-stained sEVs, were included in each experiment to define background signal and confirm antibody specificity. Data was acquired using NanoFCM Professional Suite v2.0 software and analyzed with FlowJo software (v.10.10.0).

### Imaging extracellular vesicles using dSTORM microscopy

sEVs were prepared for dSTORM imaging using the ONI EV Profiler Kit protocol. Briefly, 10^10^ of sEVs or pEVs were immobilized on chips via S3 and S4 capture molecules or on 0.1% (w/v) poly-L-lysine-coated coverslips, followed by fixation with 4% paraformaldehyde (PFA). For immunostaining, rEVs or pEVs were incubated for 2 hours at room temperature with fluorescently labeled primary antibodies against tetraspanins (CD9, CD63, CD81), diluted 1:100 in PBS containing 0.1% BSA. After PBS washing and post-fixation with 4% PFA, samples were mounted in a BCubed imaging buffer. Imaging was performed on the ONI NanoImager using three laser channels (640 nm, 561 nm, and 488 nm) with a 100× objective (50 μm × 80 μm field of view). Acquired images were analyzed using CODI software

### Isolation of EVs by an iodixanol density cushion and iodixanol density gradient (IDC+IDG)

Blood plasma sEVs isolation protocol was adapted and modified from^43^. Briefly, total of 2mL plasma was added to an ultracentrifuge tube and then 2mL of 60% iodixanol was carefully laid at the bottom of the tube (Utra-Clear, Centrifuge Tubes, product no: 344059, Beckman). The samples were then centrifuged at 100,000 × g (SW 41 Ti rotor, Beckman Coulter, USA) for 3h at 4°C. A needle was then used to aspirate 3-mL from the bottom of the tube, which included the 2-mL of iodixanol and the 1-mL of the solution on top of it, which produced a mixture of the sample in 40% iodixanol. This 3-mL solution was further fractioned on an iodixanol density gradient (IDG). Briefly, a total of 3-mL concentrate was mixed with 1-mL of 60% iodixanol and laid at the bottom of the tube, and 1-mL layers of 38%, 35%, 30%, 26%, 24%, 22%, 20% and 10% iodixanol were carefully overlaid forming a discontinuous gradient. The gradient was then centrifuged at 120,000 × g (SW 41 Ti, Beckman Coulter, USA) for 16h at 4°C. After centrifugation, 1-mL fractions were collected from the top to the bottom of the tube

### Data availability

All dPCR, nFCM fcs raw data files are available upon request

### Code availability

Access PICO BioScience’s free online PI-QUANT software for the data analysis (https://pico-bioscience.shinyapps.io/piquant/).

## Acknowledgements

We acknowledge the support of the Facility for Extracellular Vesicle Analysis and Liquid Biopsy (EV-Core), Medical Center – University of Freiburg (registered with the German Research Foundation [DFG] under RI_00612), and the financial support it received from the Faculty of Medicine – University of Freiburg (project 2025/A13-Naz).

Electron microscopy data were collected at the Cryo-EM Facility of the University of Freiburg (RRID: SCR_025860). The Titan Krios G4 cryo-TEM used for imaging was funded by Deutsche Forschungsgemeinschaft (project no. 506518771) and is operated within the Microscopy and Image Analysis Platform (MIAP), University of Freiburg.

This work was supported by German Federal Ministry for Research, Technology and Space (BMFTR, projects EV-Surf 13GW0605F; nanodiag_BW P3 03ZU1208CA), and the State Ministry for Science, Research and Arts Baden-Württemberg (MWK, project NaPeGen MWK33-7532-50/1/3).

## Author notes

These authors contributed equally: John Atanga, Pablo Sánchez-Martín.

## Use of generative artificial intelligence tools

Generative artificial intelligence (ChatGPT, OpenAI) was used exclusively for language and stylistic editing of the manuscript. All content was reviewed and validated by the authors, who take full responsibility for the work.

