## Extended data figures for "Reference-free single-vesicle profiling of small extracellular vesicles from liquid biopsies with the PICO assay"

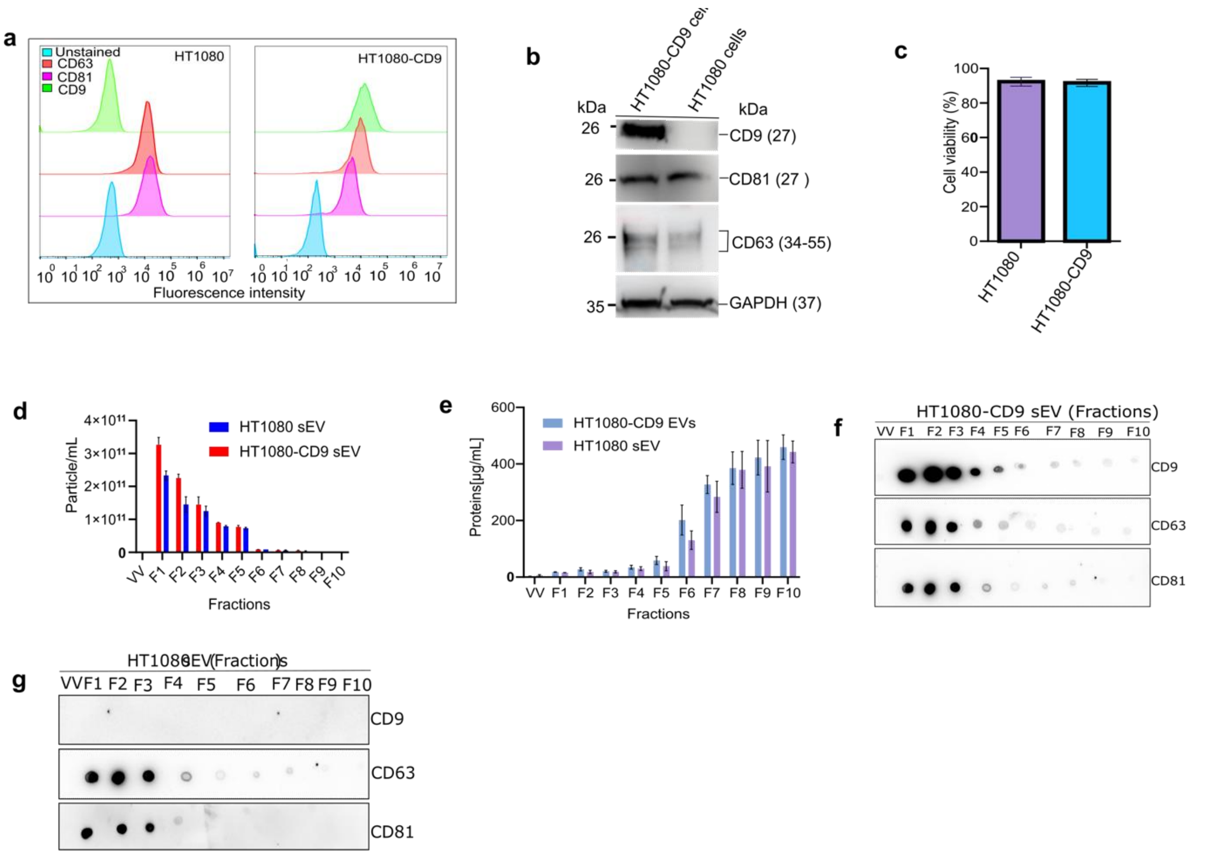


***Extended Data Fig. 1 Characterization of engineered HT1080 cell lines and sEV isolation.a,*** *Flow cytometry analysis confirming surface expression of CD9, CD63, and CD81 on HT1080 and HT1080-CD9 cells.* ***b,*** *Immunoblot analysis of CD9, CD63, and CD81 expression in parental HT1080 (CD9$^-$) and HT1080-CD9 (CD9$^+$) knock-in cell lysates.* ***c,*** *Cell viability assessment at the time of conditioned media harvest for sEV isolation. Data are presented as mean ± s.d.; n=3 biological replicates.* ***d, e,*** *Characterization of size-exclusion chromatography (SEC) fractions. NTA particle concentration* ***(d)*** *and protein concentration* ***(e)*** *across SEC fractions 1-10. Fractions 1-3, characterized by high particle count and low protein content, were selected and pooled, further concentrated for subsequent experiments.* ***f, g,*** *Dot blot analysis of SEC fractions (1-10) from HT1080-CD9* ***(f)*** *and HT1080* ***(g)*** *sEVs, probed for the indicated tetraspanins.*

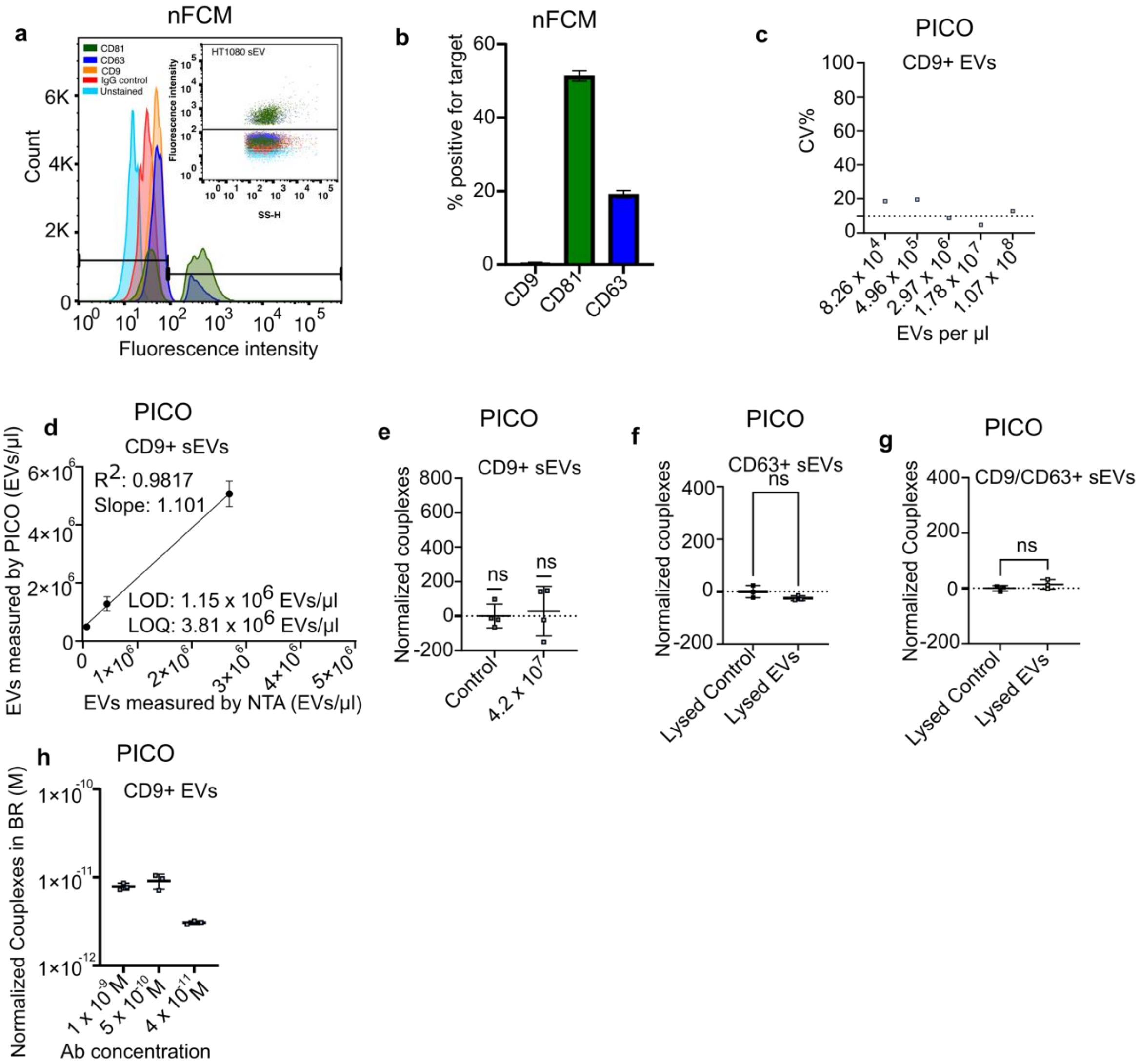


**Extended Data Fig. 2 Analytical validation of the single-marker PICO assay.*a, b,*** *Single-vesicle analysis of parental HT1080 ($CD9^-$) sEVs by nFCM. Representative scatter plots **(a)** and quantification **(b)** of CD9, CD81, and CD63 positivity, confirming the absence of CD9 signal.* ***c,*** *Inter-assay precision of the PICO single-marker assay. Coefficient of variation (CV) for technical and biological replicates using HT1080-CD9 sEVs and CD9 as detection marker. Dashed line indicates 10% CV threshold.* ***d,*** *Determination of the limit of detection (LOD, 1.15 × $10^6$ sEV/µL) and limit of quantification (LOQ, 3.81 × $10^6$ sEV/µL) for CD9+ sEVs using the single-marker PICO assay.* ***e,*** *CD9 detection on HT1080-CD9 sEVs, no signals were detected indicating high specificity of PICO assay, since this sEVs lack CD9 expression.* ***f, g,*** *Specificity control for intact sEV detection. PICO single-marker (CD63, **f**) or double-marker (CD9/CD63, **g**) assay performed on lysed HT1080-CD9 sEVs. Data are presented as mean ± s.d.; n=2 biological replicates.* ***h,*** *Antibody titration for saturation determination. Corected couplex counts for CD9+ sEVs were measured across a range of antibody concentrations.*

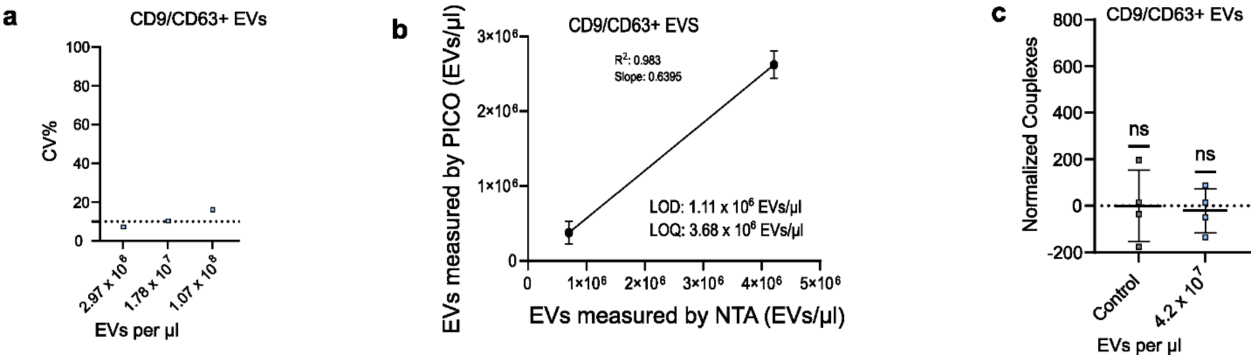


***Extended Data Fig. 3 Analytical validation of the double-marker PICO assay.***

***a,*** *Inter-assay precision of the PICO double-marker assay. Coefficient of variation (CV) for technical and biological replicates across CD9/CD63 detection.* ***b,*** *Determination of the limit of detection (LOD, 1.11 × $10^6$ sEV/µL) and limit of quantification (LOQ, 3.68 × $10^6$ sEV/µL) for CD9/CD63+ sEVs using the double-marker PICO assay.* ***c,*** *Specificity control for double-marker detection. PICO double-marker assay (CD9/CD63) performed on HT1080 ($CD9^-$) sEVs. Normalized couplex counts show no significant signal above the antibody binding*

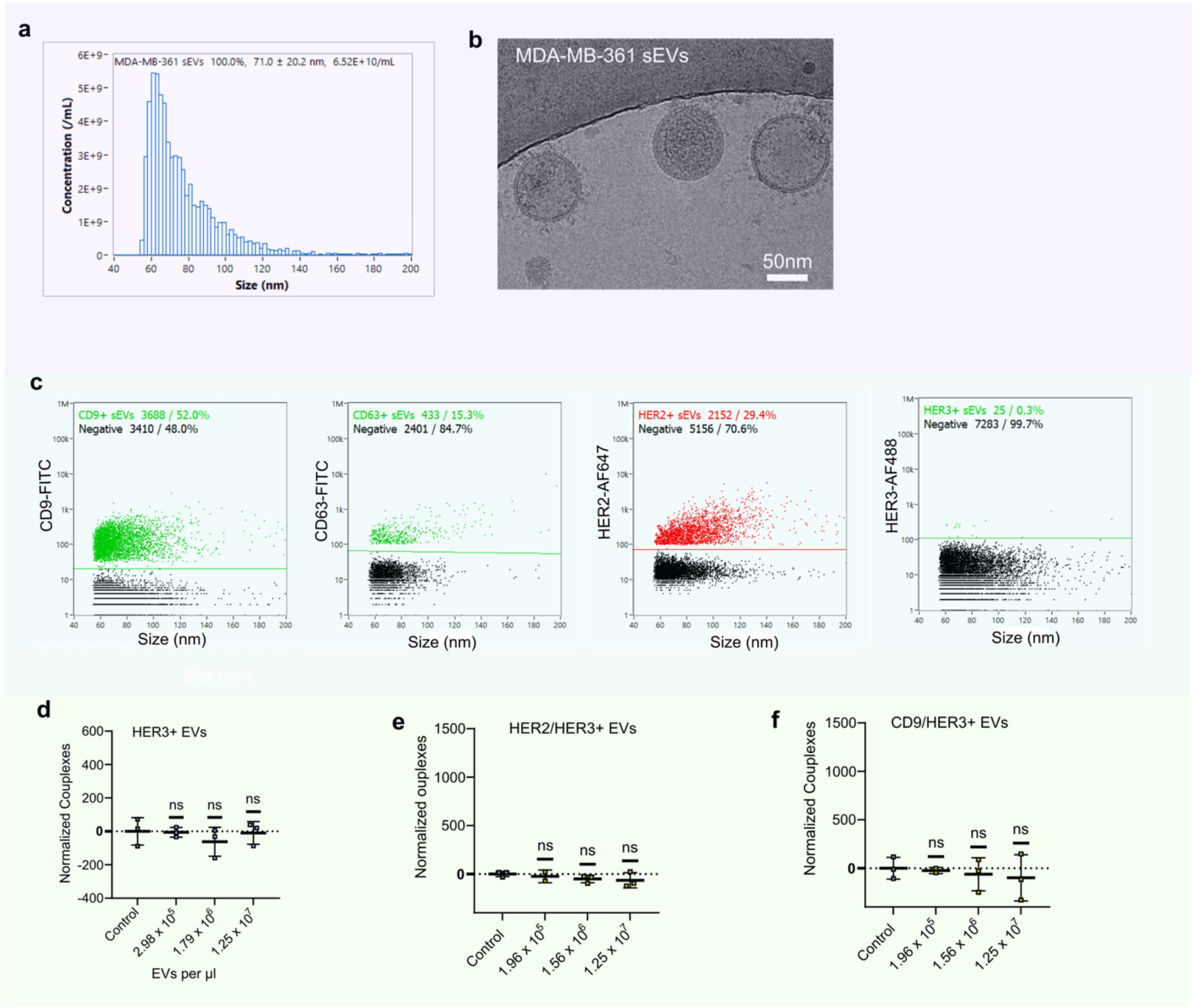


*control (ABC).*

**Extended Data Fig. 4 Characterization of MDA-MB-361-derived sEVs and PICO specificity for tumor-relevant markers. a,** Nanoparticle tracking analysis (NTA) of sEVs derived from the HER2+ breast cancer cell line MDA-MB-361. Representative size distribution profile. **b,** Representative cryo-electron microscopy (cryo-EM) image of purified MDA-MB-361-derived sEVs, showing intact bilayered membranes. Scale bar, 50 nm. **c,** Single-vesicle analysis of MDA-MB-361-derived sEVs by nFCM. Quantification of CD9, CD63, HER2, HER3 positivity, no signals were detected for HER3 confirming its absence on these sEVs. **d,** PICO single-marker dilution series for HER3 on MDA-MB-361-derived sEVs, showing no detectable signal across a range of sEV concentrations. **e, f,** PICO double-marker assay for HER2/HER3 **(e)** and CD9/HER3 **(f)** on MDA-MB-361-derived sEVs, confirming the absence of detectable Couplexes for combinations involving HER3.

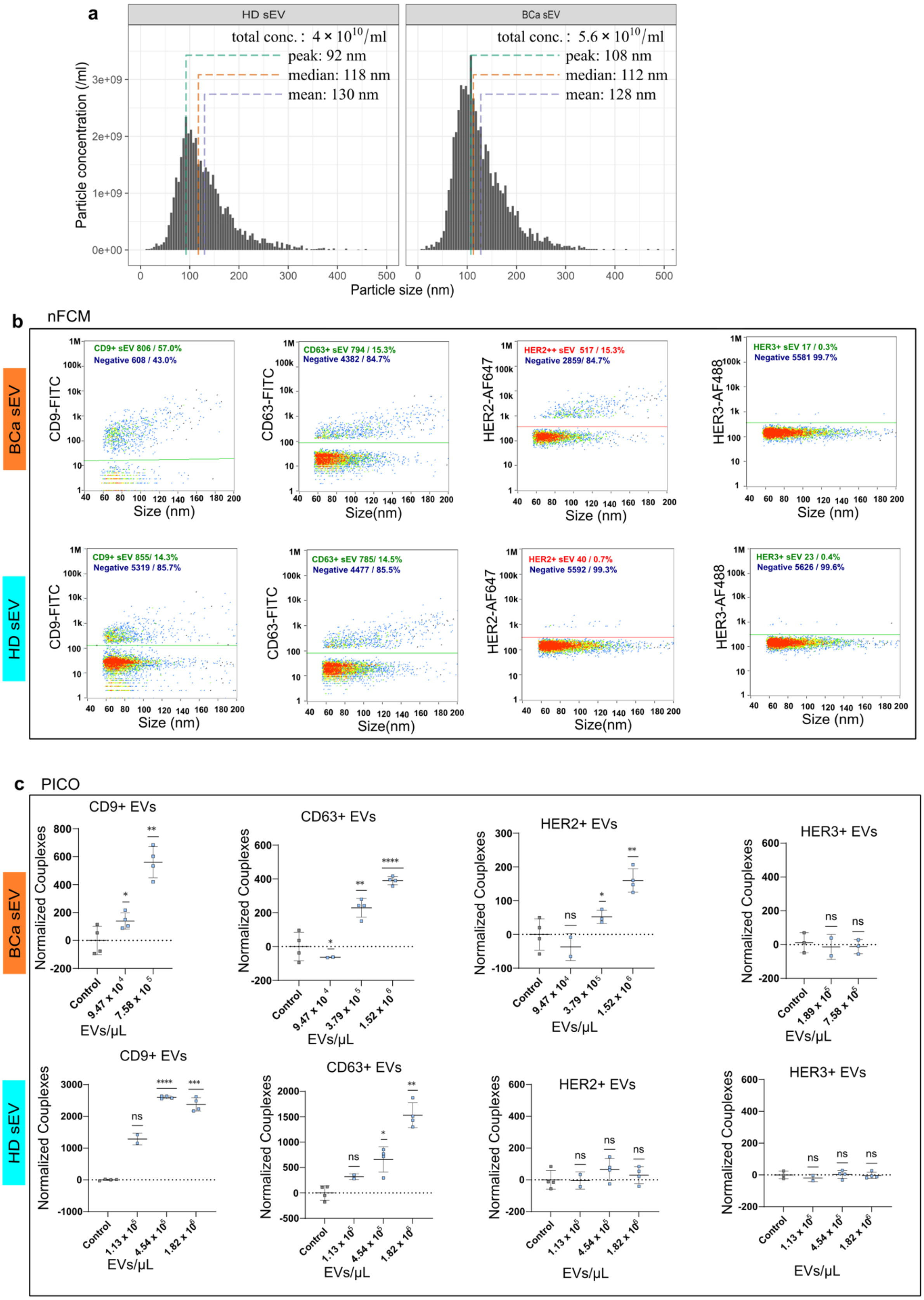
a
HD sEV
BCa sEV
total conc.: 4 × 10^10/ml
peak: 92 nm
median: 118 nm
mean: 130 nm
total conc.: 5.6 × 10^10/ml
peak: 108 nm
median: 112 nm
mean: 128 nm
Particle concentration (/ml)
Particle size (nm)
b nFCM
BCa sEV
HD sEV
CD9+ sEV 806 / 57.0%
Negative 608 / 43.0%
CD63+ sEV 794 / 15.3%
Negative 4382 / 84.7%
HER2++ sEV 517 / 15.3%
Negative 2859/ 84.7%
HER3+ sEV 17 / 0.3%
Negative 5581 99.7%
CD9+ sEV 855/ 14.3%
Negative 5319 / 85.7%
CD63+ sEV 785/ 14.5%
Negative 4477 / 85.5%
HER2+ sEV 40 / 0.7%
Negative 5592 / 99.3%
HER3+ sEV 23 / 0.4%
Negative 5626 / 99.6%
CD9-FITC
CD63-FITC
HER2-AF647
HER3-AF488
Size (nm)
c PICO
CD9+ EVs
CD63+ EVs
HER2+ EVs
HER3+ EVs
Normalized Couplexes
EVs/μL
Control
9.47 x 10^4
7.58 x 10^5
3.79 x 10^5
1.52 x 10^6
1.89 x 10^5
1.13 x 10^6
4.54 x 10^6
1.82 x 10^6
ns

**Extended Data Fig. 5 Characterization of plasma-derived sEVs from breast cancer patients and healthy donors with single-marker detection. a,** Size distribution analysis by NTA of sEVs isolated from breast cancer patients (n=4) and matched healthy donors (n=4); **b, c,** Analysis of CD9, CD63, HER2 and HER3 expression. nFCM upper panel represent the breast cancer samples indicated as (BCa) and lower panel represent healthy individual samples indicated as (HD) **(b)**. PICO single-marker assay, upper and lower panel represent BCa and HD respectively **(c)** confirm the absence of HER3+ sEVs in plasma samples from both breast cancer patients and healthy donors. HER2+ sEVs was specifically detected in breast cancer patients.

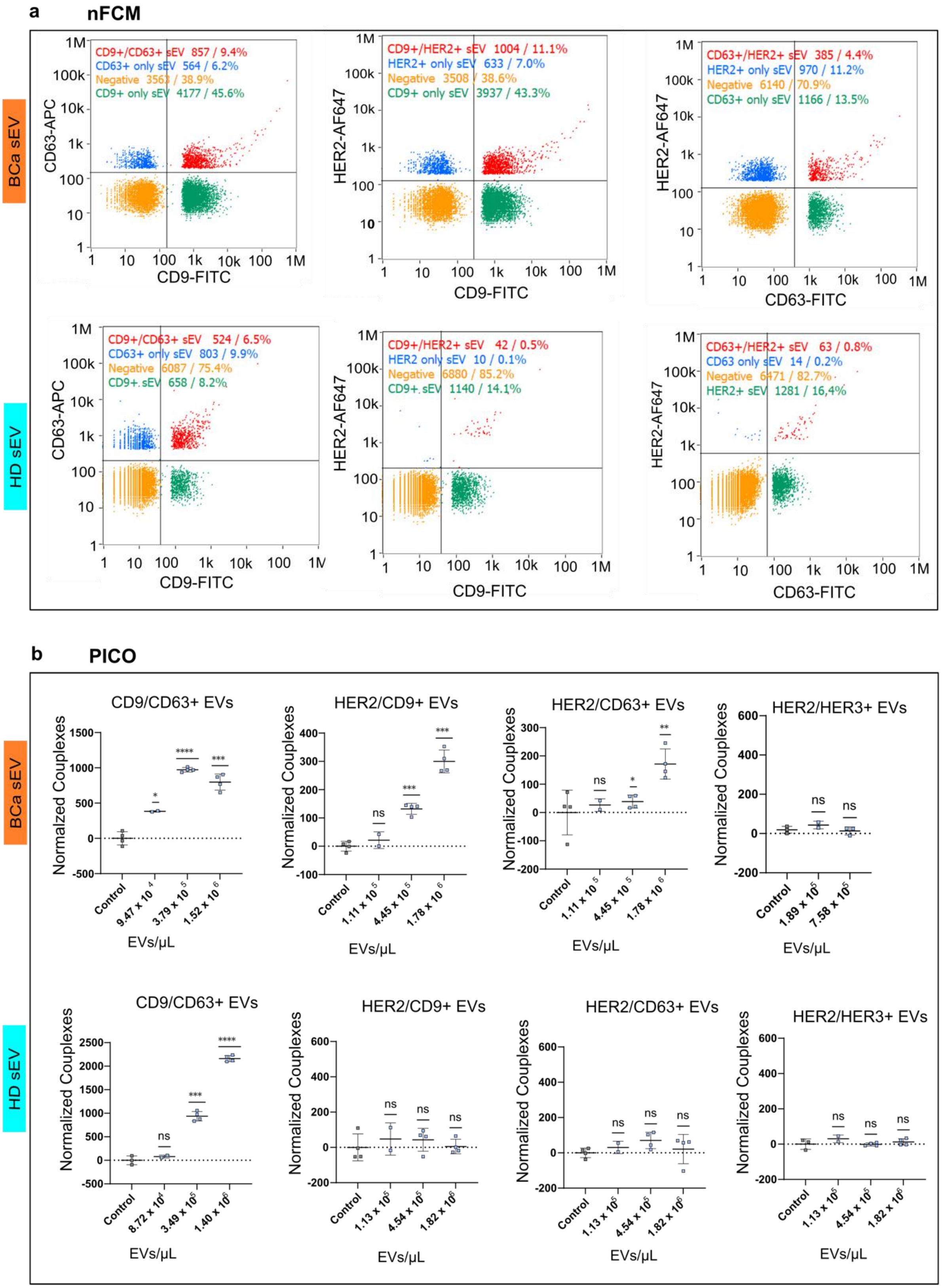
a
nFCM
BCa sEV
HD sEV
CD9+/CD63+ sEV 857 / 9.4%
CD63+ only sEV 564 / 6.2%
Negative 3563 / 38.9%
CD9+ only sEV 4177 / 45.6%
CD9+/HER2+ sEV 1004 / 11.1%
HER2+ only sEV 633 / 7.0%
Negative 3508 / 38.6%
CD9+ only sEV 3937 / 43.3%
CD63+/HER2+ sEV 385 / 4.4%
HER2+ only sEV 970 / 11.2%
Negative 6140 / 70.9%
CD63+ only sEV 1166 / 13.5%
CD9+/CD63+ sEV 524 / 6.5%
CD63+ only sEV 803 / 9.9%
Negative 6087 / 75.4%
CD9+ sEV 658 / 8.2%
CD9+/HER2+ sEV 42 / 0.5%
HER2 only sEV 10 / 0.1%
Negative 6880 / 85.2%
CD9+ sEV 1140 / 14.1%
CD63+/HER2+ sEV 63 / 0.8%
CD63 only sEV 14 / 0.2%
Negative 6471 / 82.7%
HER2+ sEV 1281 / 16,4%
CD63-APC
HER2-AF647
CD9-FITC
CD63-FITC
b
PICO
CD9/CD63+ EVs
HER2/CD9+ EVs
HER2/CD63+ EVs
HER2/HER3+ EVs
Normalized Couplexes
EVs/µL
Control
9.47 x 10 4
3.79 x 10 5
1.52 x 10 6
1.11 x 10 5
4.45 x 10 5
1.78 x 10 6
1.89 x 10 5
7.58 x 10 5
8.72 x 10 4
3.49 x 10 5
1.40 x 10 6
1.13 x 10 5
4.54 x 10 5
1.82 x 10 6
ns

**Extended Data Fig. 6 Specificity of double-marker PICO detection in clinical plasma samples. a,** Representative scatter plots from nFCM analysis showing the percentage of sEV subpopulations co-expressing CD9/CD63, HER2/CD9, and HER2/CD63 in breast cancer patient and healthy donor plasma samples. **b,** Specificity of the PICO double-marker assay. In healthy donor samples, even if CD9 or CD63 antibodies bind to sEVs, the absence of HER2 binding prevents the formation of a detectable Couplex signal, resulting in no counts for HER2-containing pairs.
